# Establishment of *Drosophila* Intestinal Cell Lines

**DOI:** 10.1101/2025.05.27.656384

**Authors:** Arthur Luhur, Daniel Mariyappa, Peter Bohall, Laura Multini, Morgan Elkins, Andrew C. Zelhof

**Affiliations:** Drosophila Genomics Resource Center Biology Department Indiana University Bloomington, IN 47405

**Keywords:** *Drosophila*, intestine, cell line, spheroids, transcriptomics

## Abstract

The *Drosophila* intestine is a powerful model for stem cell dynamics and epithelial biology, yet no intestinal derived cell lines have been available until now. Here, we describe the establishment of *Drosophila* intestinal cell lines. The cell lines were derived from the embryonic intestine through specific Ras^V12^ expression and show continuous proliferation and the ability to be frozen and re-thawed. Each derived cell line exhibited morphological and cellular heterogeneity. Single-cell RNA sequencing confirmed their intestinal origin and cell populations with unique enriched signaling pathways. In addition, L15, one of the three lines, formed 3D spheroids that displayed epithelial polarity. Together, these lines provide an additional resource for studying intestinal development, epithelial organization, and pest management.

## Introduction

Insect cell lines are invaluable complementary tools for dissecting fundamental biological processes, unraveling host-pathogen interactions, and driving biotechnological advancements^1–3^. Among these, *Drosophila melanogaster*, with its rich genetic toolkit and well-characterized developmental biology, stands as a critical model organism for unraveling the complexities of cellular and molecular processes. However, despite the vast utility of *Drosophila* cell cultures in elucidating diverse biological processes, the establishment of intestine-derived cell lines from this model organism has remained a significant gap in the research landscape.

The intestine represents a pivotal organ in insects, serving critical roles in nutrient digestion, absorption, xenobiotic metabolism, and immune defense^4,5^. Studies on intestine physiology and pathology are fundamental for understanding insect biology and for developing novel approaches in insect pest management. While *Drosophila* intestine tissues have been extensively studied *in vivo*, the lack of corresponding cell lines has limited the scalability of investigations into intestine-specific processes.

Although numerous *Drosophila* cell lines have been established and characterized— such as the hemocyte-derived S2 and Kc167 lines ^6,7^, and more recently, epithelial, neuronal, and glial lines from late-stage embryos^8^, none have recapitulated the intestinal tissue, therefore leaving a void for modeling gut functions *in vitro*.

Beyond developmental biology, insect intestinal cell lines enable the study of various gut toxins, including those from *Bacillus thuringiensis* (Cry and Cyt toxins)^9^, *Photorhabdus* (Toxin complex and PirAB toxins)^10^, plant-derived compounds such as insecticidal cyclotides^11^, spider or scorpion toxins that target gut functions^12,13^. These systems are also well suited for exploring insecticide detoxification. For instance, the insect ATP-Binding Cassette (ABC) transporters in the insect gut are critical for expelling xenobiotics and likely play a significant role in the development of insecticide resistance^14,15^. In addition, *Drosophila* intestinal cell lines with site-specific transgene insertion sites could provide a robust platform for investigating the cellular and molecular mechanisms by which insecticidal toxins affect the gut. For instance, the cell line could facilitate the functional characterization of specific heterologous ABC transporters, including orthologs derived from other insect species. Moreover, these cell lines could be used as high-throughput screening platforms for pest control candidates.

Here, we describe the establishment and characterization of intestinal-derived cell lines from *Drosophila melanogaster,* using the UAS-GAL4 targeted expression of activated oncogene Ras85D^8,16,17^ in the late embryonic intestine. The derived lines also carry a chromosomal *attP* landing site for precise transgene integrations. The characterization of the derived lines recapitulates aspects of intestine biology including transcriptomic profiles mapping to intestinal tissue, known cell type markers, and epithelial polarity.

One of these lines was also able to self-organize to form spheroids with apical-basal polarity, demonstrating its propensity to form 3D structures. We anticipate these lines will support further investigations into insect intestinal physiology and serve as scalable *in vitro* tools for the research community.

## Results

### Establishing continuous cell cultures using embryonic gut-specific CG10116- GAL4 driver

Embryonic tissues are highly plastic and proliferative, making them suitable for establishing intestinal cell lines. Embryos are also easy to surface sterilize, which helps avoid the contamination issues that make intestinal cultures from larval or adult stages more challenging to obtain. To establish primary cultures that favor the proliferation and survival of cells from the intestinal lineage, we utilized expression of oncogenic Ras85D (Ras^V^^12^)^8,17^. To specifically target the embryonic intestinal expression of oncogenic Ras^V^^12^, we utilized CG10116-GAL4. CG10116-GAL4 was previously shown to direct the expression of a UAS-GFP reporter in most cells in the larval and adult intestines, and in a small number of cells in the larval Malpighian tubules^18^. Single cell transcriptomic data from the adult gut indicated that CG10116 is expressed and enriched in the differentiated enterocytes (ECs) relative to the other cell types^19^. In the developing embryonic gut, GFP expression driven by CG10116-GAL4 was first faintly detected in the posterior midgut between embryonic stages 11 and 12, at the point of the posterior midgut folding ventrally (Supplementary Fig. 1). At this stage, no GFP signal was observed in the anterior midgut. By stage 12, faint GFP expression appeared in the anterior midgut, while the posterior midgut showed a stronger signal. At stage 13, GFP signal was robust in the developing midgut chamber, particularly on the ventral side.

From stages 13 to 17, GFP expression persisted in the midgut and was also likely present in some cells of the Malpighian tubules as midgut constrictions, chambers and coils developed. Throughout embryogenesis, GFP was not detected in the foregut or hindgut (Supplementary Fig. 1).

Next, we tested whether expression Ras^V12^ affected the development of the intestine and viability of the animal. Although constitutively active Ras^V12^ was expressed in the midgut, the gross morphology of the embryonic midgut was comparable to that of wild- type embryos (Fig. 1A – B). In addition, animals expressing Ras^V12^ in their intestine progressed through development, survived the larval stages, and became fertile adults. GFP expression, serving as a proxy for Ras^V12^ expression, was also observed in the adult intestines. Ras^V12^-expressing adult intestines displayed gross morphology similar to wild-type intestines (Fig. 1C–D). Together with the data from embryonic Ras^V12^ expression, these results support the feasibility of using embryos expressing from Ras^V12^ in the midgut as a source for establishing continuous intestinal cell lines.

**Figure 1.**
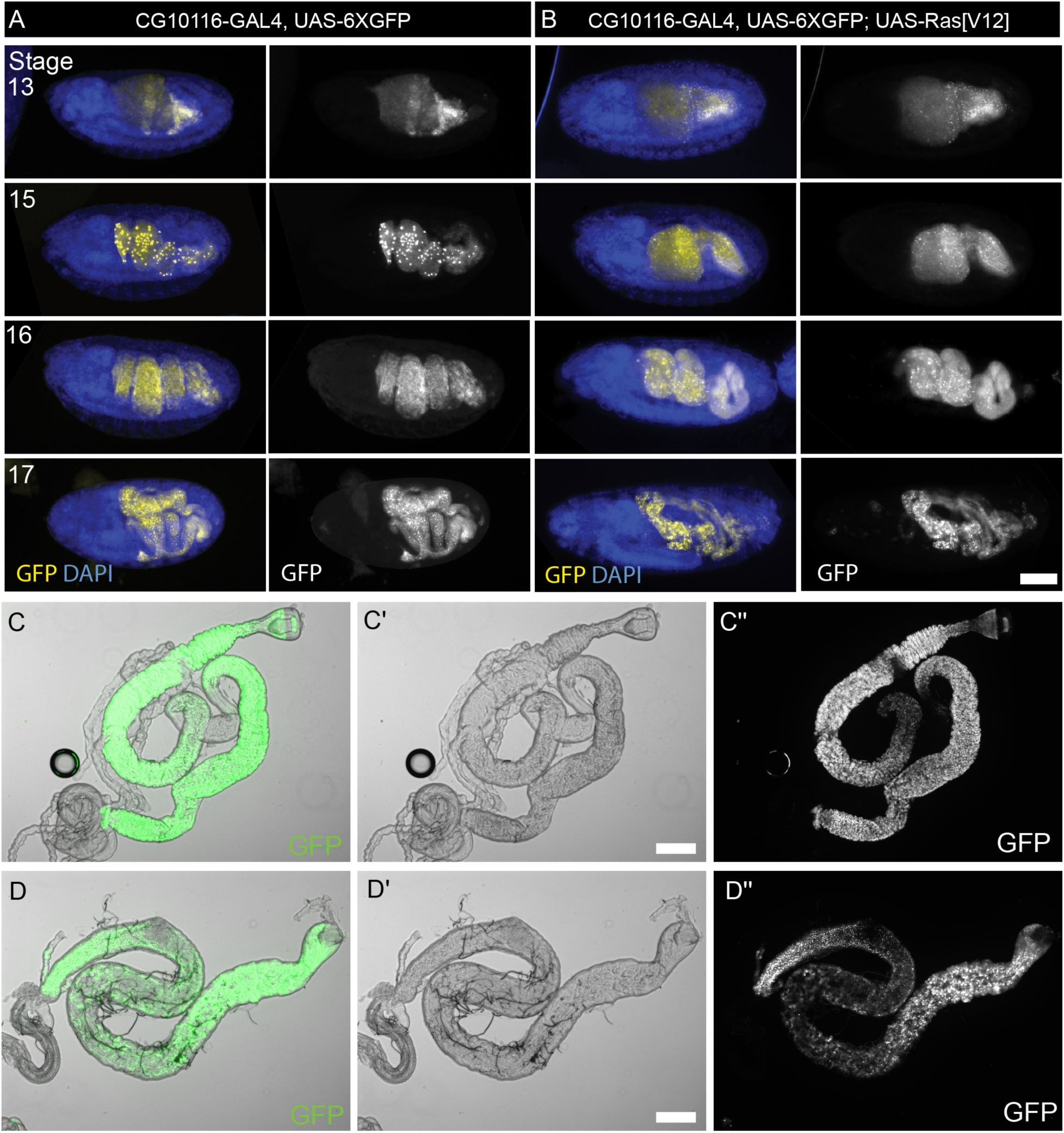
CG10116-GAL4 drives intestinal-specific GFP expression in embryos and adults. Micrographs of Stage 13 to Stage 17 wildtype (A) or Ras^V12^ expressing (B) embryos. To the left of each panel are merged images with DAPI and GFP (yellow) and the images to the right are GFP channel only. Micrographs of the GFP channel highlighting intestinal morphology of the embryos expressing Ras^V12^ in the intestinal cells are displayed to the right. GFP expression driven by CG10116-GAL4 was also characterized in either control (C) or Ras^V12^ (D) adult male guts. The respective brightfield (C’,D’) and GFP channel (C’’, D’’) are also shown. Scale bars = 100 µm.

Given the specificity of CG10116-GAL4 and the absence of lethality associated with Ras^V12^ expression, we generated embryos from a genetic cross in which the offspring were co-expressing Ras^V12^ and GFP. To enable phiC31 integrase-mediated Recombinase-Mediated Cassette Exchange (RMCE)^20^, the embryos were also designed to carry a single genomic cassette containing inverted *attP* sites flanking a white transgene inserted at the *tara* locus on the third chromosome^21^ (details in Methods and Fig. 2). We derived twenty independent primary cultures from embryos aged for 18-23 hours post oviposition at 25°C which are enriched for Stages 16-17 embryos. (Fig. 2B). Seven of the twenty progressed to become continuous growing cultures over the period four to five weeks. In the end, we cryopreserved five of the seven cultures. Here, we describe the physical characteristics of the three of the five continuous cell lines: CG10116-L10 (L10), CG10116-L15 (L15) and CG10116-L18 (L18).

**Figure 2.**
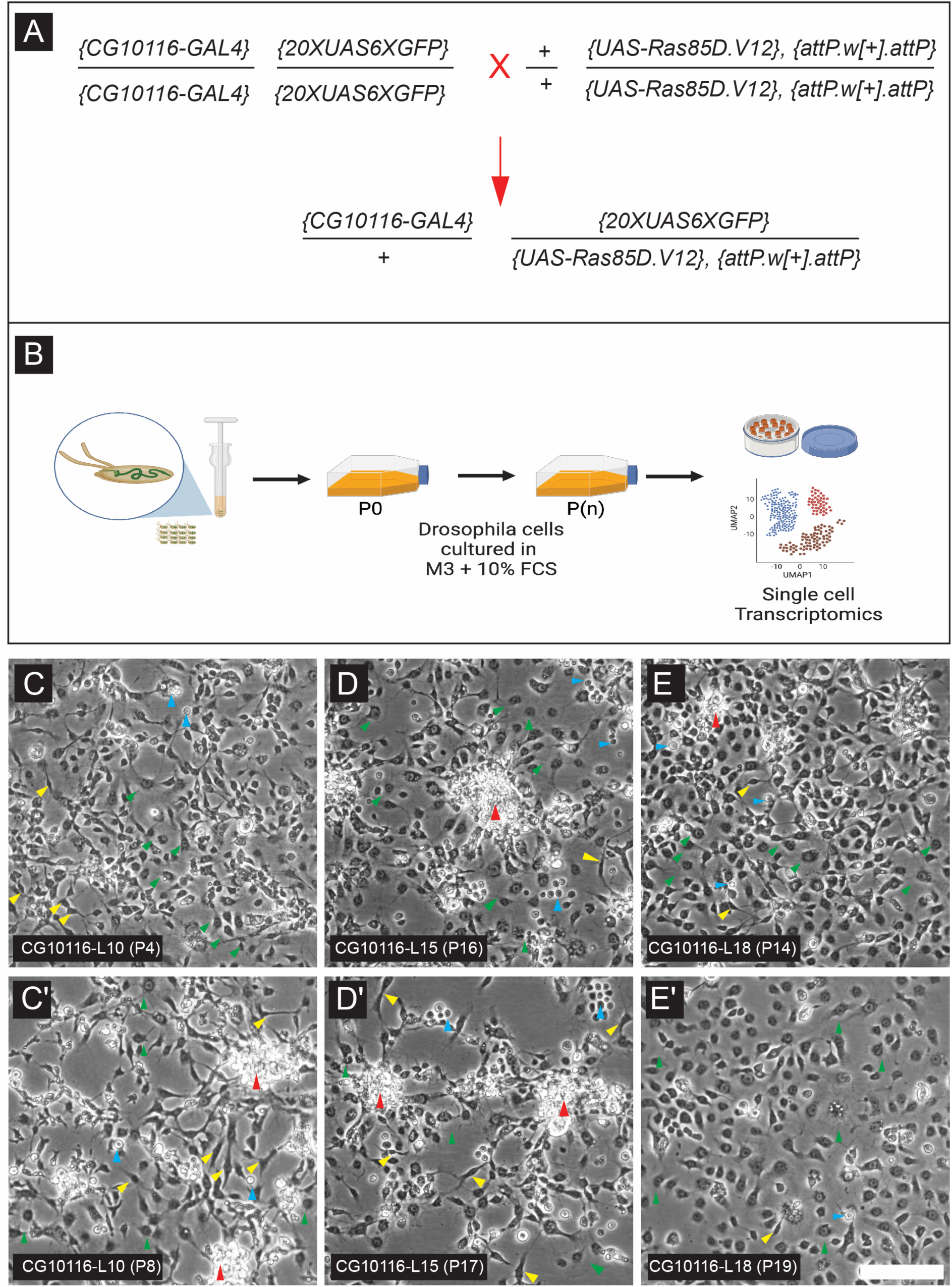
Strategy to create and morphology of embryonic intestine derived cell lines. (A)The schematic for the genetic cross to generate embryos carrying the CG10166-GAL4 driver together with UAS-6XGFP, constitutively active Ras85D (UAS- Ras^V12^) and a copy of an attP insertion site on the third chromosome. (*B)* Established cell lines were cryopreserved, analyzed also subjected to single cell transcriptomic analysis. (C-C’) Micrographs of L10 at passage 4 and passage 8, respectively. (D-D’) Micrographs of L15 at passage 15 and passage 17, respectively. (E-E’) Micrographs of L18 at passage 14 and passage 19, respectively. Scale bar = 100 µm. Red arrowheads indicate cell aggregates. Green arrowheads indicate epithelial-like cells. Blue arrowheads indicate round cells. Yellow arrowheads indicate spindle-shaped cells.

## Morphological and growth characteristics of intestinal cell lines

All three continuous cell lines L10, L15 and L18 were derived from embryos having the same genotype, and contain cells that exhibit unique and mixed morphological features. The shapes include spindle-shaped cells, epithelial-like cells, small round cells and cell clumps that indicated a lack of contact inhibition (Fig. 2C-E, Supplementary Fig. 2A-C). Not all cells expressed GFP, especially in L15 (Supplementary Fig. 2B). The growth profiles of these lines were comparable (Supplementary Fig. 2D-F). L10 and L15 reached confluent density of 3 million cells/mL, while L18 reached its confluent density at 1 million cells/mL. Doubling time ranged from 50-70 hours (Supplementary Fig. 2J).

We also measured the chromosomal content via cell sorting. L10 and L15 have comparable DNA content profile, with cells in the 2n, 4n, and 8n phases (Supplementary Fig. 2G-H). In contrast, L18 displayed two main peaks at 2n and 4n (Supplementary Fig. 2I). The mixed ploidy observed in these cell lines may correspond to known population of cells in the *Drosophila* midgut. The diploid (2n) population may represent the larval Adult Midgut Progenitor (AMP) cells or the secretory enteroendocrine (EE) cells^22–25^, while the 4n population may represent pre-mitotic tetraploid cells, cells undergoing endoreplication, or cells that are transitioning toward polyploid ECs, as indicated by the presence of cells with 8n ploidy^26^.

### Single cell profiling confirms heterogeneity of the intestinal cell lines

The process of establishing a continuous cell culture provides a unique opportunity to capture the diverse gene expression states that result from cells adapting to continuous *in vitro* growth and cell division outside their physiological niches. To gain insight into the polyclonal nature of the three independently derived lines, we performed single cell RNA-sequencing (scRNA-seq). Our goal was to characterize the types of cell types present in each line. scRNA-seq enables this resolution by identifying distinct cell types and offering a detailed snapshot of cellular heterogeneity. We dissociated L10 (Passage 4), L15 (Passage 18), and L18 (Passage 16) using trypsin to obtain single-cell suspensions. Consistent with observed differences in morphology and ploidy, UMAP projections of the transcriptomes revealed the polyclonal nature in all three lines. Seurat clustering identified 18, 9, and 12 distinct clusters in L10, L15, and L18, respectively (Fig. 3).

**Figure 3.**
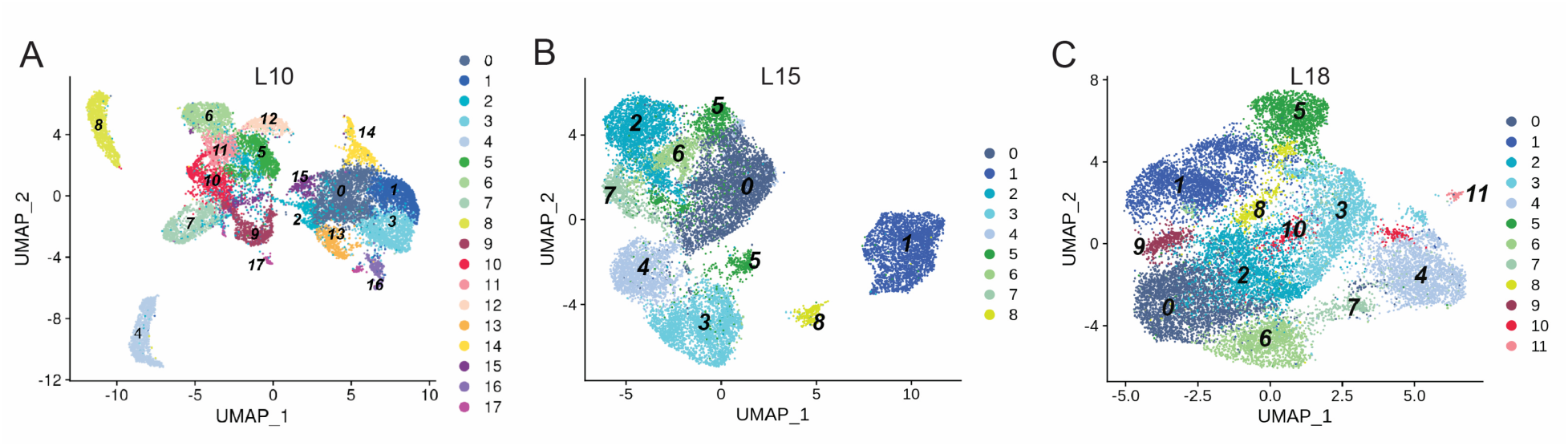
UMAP of the intestinal-derived Drosophila cell lines for L10, L15 and L18 samples by single cell transcriptomics. (A) Uniform Manifold Approximation and Projection (UMAP) of the 18 clusters identified from 15,076 L10 cells at passage 5. (B) UMAP of the 9 clusters identified from 16,472 L15 cells at passage 18. (C) UMAP of the 12 clusters identified from 17,695 L18 cells at passage 16.

To obtain an insight into what these cells resemble molecularly, we used the Drosophila RNAi Screening Center’s single-cell Database (DRscDB) tool^27^ to compare cluster enriched genes against the Fly Cell Atlas (FCA) single-cell RNAseq genesets^28^. Using the whole-body FCA geneset, we found many clusters from the three lines significantly expressed gene signatures characteristic of the gut tissue (Table 1). Given this result, we reasoned that a more targeted comparison to the intestine-specific FCA dataset would provide higher-resolution insights. This tissue-focused dataset captures the full diversity of intestinal cell types and differentiation states, allowing us to assess whether our cultured cells align with physiological cell populations from the adult gut. Indeed, we found that six out of the nine clusters in L15 were significantly enriched for intestinal stem cell (ISC) markers, with the remaining clusters expressing genes characteristic to the adult enteroblast (EB), cardia and the malpighian tubule (Table 2). In L18, five out of twelve clusters significantly expressed genes for ISC markers, while the remaining clusters expressed genes characteristic of other cell types in the gut including the EBs, and differentiating ECs. In L10, eleven out of eighteen clusters expressed genes characteristic of the ISCs, with the remaining clusters expressed genes characteristic of other cell types of the adult gut, including EBs, differentiating EC, cardia, and EEs (Table 2).

**Table 1.** DRscDB analysis for L15, L10, and L18, against the whole body_dm_FCAvs2_10x geneset.

| **L15**<br>**Cluster** | **Cell cluster scRNAseq** | **p value** | **Tissue/Cell** | **Rank** |
| --- | --- | --- | --- | --- |
| 0 | intestinal stem cell | 1.33E-17 | intestine | 2 |
| 1 | midgut | 1.22E-23 | intestine | 2 |
| 2 | intestinal stem cell | 3.12E-19 | intestine | 1 |
| 3 | intestinal stem cell | 4.74E-12 | intestine | 2 |
| 4 | intestinal stem cell | 4.34E-06 | intestine | 3 |
| 5 | intestinal stem cell | 1.25E-21 | intestine | 3 |
| 6 | intestinal stem cell | 5.92E-11 | intestine | 1 |
| 7 | adult hindgut | 7.95E-04 | intestine | 10 |
| 8 | adult hindgut | 1.34E-12 | intestine | 3 |

| **L18**<br>**Cluster** | **Cell cluster scRNAseq** | **pvalue** | **Tissue/Cell** | **Rank** |
| --- | --- | --- | --- | --- |
| 0 | midgut | 1.00E-25 | Gut | 1 |
| 1 | adult tracheal cell | 4.75E-05 | Trachea | 3 |
| 2 | intestinal stem cell | 1.12E-13 | Gut | 7 |
| 3 | intestinal stem cell | 6.71E-08 | Gut | 11 |
| 4 | midgut | 4.61E-17 | Gut | 6 |
| 5 | adult midgut enterocyte | 3.82E-04 | Gut | 4 |
| 6 | intestinal stem cell | 5.80E-03 | Gut | 12 |
| 7 | midgut | 6.57E-03 | Gut | 5 |
| 8 | adult midgut-hindgut hybrid zone | 1.21E-02 | Gut | 6 |
| 9 | intestinal stem cell | 2.10E-13 | Gut | 7 |
| 10 | intestinal stem cell | 6.83E-25 | Gut | 2 |
| 11 | intestinal stem cell | 3.73E-15 | Gut | 7 |

| **L10**<br>**Cluster** | **Cell cluster scRNAseq** | **pvalue** | **Tissue/Cell** | **Rank** |
| --- | --- | --- | --- | --- |
| 0 | adult hindgut | 3.75E-02 | Gut | 15 |
| 1 | intestinal stem cell | 2.57E-09 | Gut | 10 |
| 2 | midgut | 1.97E-03 | Gut | 4 |
| 3 | intestinal stem cell | 3.27E-15 | Gut | 1 |
| 4 | midgut | 1.00E-25 | Gut | 4 |
| 5 | midgut | 1.00E-25 | Gut | 1 |
| 6 | midgut | 4.03E-11 | Gut | 3 |
| 7 | cardia | 6.13E-15 | Gut | 44 |
| 8 | midgut | 1.00E-25 | Gut | 1 |
| 9 | intestinal stem cell | 1.34E-15 | Gut | 9 |
| 10 | intestinal stem cell | 5.11E-14 | Gut | 9 |
| 11 | midgut | 2.16E-21 | Gut | 2 |
| 12 | intestinal stem cell | 2.93E-07 | Gut | 11 |
| 13 | intestinal stem cell | 2.06E-03 | Gut | 12 |
| 14 | intestinal stem cell | 3.53E-08 | Gut | 25 |
| 15 | cardia | 4.53E-04 | Gut | 2 |
| 16 | intestinal stem cell | 4.60E-13 | Gut | 6 |
| 17 | intestinal stem cell | 6.85E-13 | Gut | 9 |

**Table 2.** DRscDB analysis for L15, L10, and L18, against the intestine_dm_FCAvs2_10x geneset.

| **L15**<br>**Cluster** | **Cell cluster scRNAseq** | **p value** | **Tissue/Cell** | **Rank** |
| --- | --- | --- | --- | --- |
| 0 | intestinal stem cell | 1.73E-16 | intestine | 1 |
| 1 | intestinal stem cell | 1.10E-04 | intestine | 1 |
| 2 | intestinal stem cell | 1.00E-25 | intestine | 1 |
| 3 | enteroblast | 1.06E-13 | intestine | 1 |
| 4 | intestinal stem cell | 6.26E-08 | intestine | 1 |
| 5 | intestinal stem cell | 1.00E-25 | intestine | 1 |
| 6 | intestinal stem cell | 5.92E-12 | intestine | 1 |
| 7 | cardia | 2.94E-05 | intestine | 1 |
| 8 | adult malpighian tubule | 7.68E-03 | intestine | 1 |

| **L18**<br>**Cluster** | **Cell cluster scRNAseq** | **pvalue** | **Tissue/Cell** | **Rank** |
| --- | --- | --- | --- | --- |
| 0 | intestinal stem cell | 6.23E-16 | intestine | 1 |
| 1 | muscle cell | 4.75E-05 | intestine | 1 |
| 2 | intestinal stem cell | 1.00E-25 | intestine | 1 |
| 3 | enteroendocrine cell | 1.40E-06 | intestine | 1 |
| 4 | enteroblast | 1.99E-25 | intestine | 1 |
| 5 | adult midgut enterocyte | 2.37E-17 | intestine | 1 |
| 6 | intestinal stem cell | 6.48E-16 | intestine | 1 |
| 7 | adult differentiating EC | 4.16E-02 | intestine | 1 |
| 8 | adult midgut enterocyte | 2.58E-08 | intestine | 1 |
| 9 | intestinal stem cell | 1.00E-25 | intestine | 1 |
| 10 | intestinal stem cell | 5.75E-20 | intestine | 1 |
| 11 | enteroblast | 3.70E-12 | intestine | 1 |

| **L10**<br>**Cluster** | **Cell cluster scRNAseq** | **pvalue** | **Tissue/Cell** | **Rank** |
| --- | --- | --- | --- | --- |
| 0 | adult midgut enterocyte | 5.65E-08 | intestine | 1 |
| 1 | intestinal stem cell | 1.44E-17 | intestine | 1 |
| 2 | enteroblast | 1.97E-03 | intestine | 1 |
| 3 | intestinal stem cell | 1.41E-24 | intestine | 1 |
| 4 | intestinal stem cell | 1.18E-16 | intestine | 1 |
| 5 | intestinal stem cell | 9.56E-24 | intestine | 1 |
| 6 | intestinal stem cell | 1.02E-12 | intestine | 1 |
| 7 | cardia | 2.40E-13 | intestine | 1 |
| 8 | intestinal stem cell | 4.76E-18 | intestine | 1 |
| 9 | intestinal stem cell | 1.00E-25 | intestine | 1 |
| 10 | adult differentiating enterocyte | 1.16E-10 | intestine | 1 |
| 11 | intestinal stem cell | 6.28E-25 | intestine | 1 |
| 12 | enteroblast | 3.13E-14 | intestine | 1 |
| 13 | intestinal stem cell | 2.25E-09 | intestine | 1 |
| 14 | enteroendocrine cell | 1.00E-25 | intestine | 1 |
| 15 | enteroblast | 4.53E-04 | intestine | 1 |
| 16 | intestinal stem cell | 1.00E-25 | intestine | 1 |
| 17 | intestinal stem cell | 1.30E-19 | intestine | 1 |

We also reasoned that analyzing specific transcription factors known to be important for intestinal development could provide additional support for the intestinal origin for the L15, L10, and L18 cell lines. For example, in L15, the expression of *kayak* (*Drosophila* Fos-related antigen), a key regulator of intestinal development and function^29^, was highly expressed in multiple clusters (Fig. 4A–G). *Delta* and *Enhancer of Split E(spl)* transcripts, whose protein products are markers for adult intestinal progenitor cells, were also enriched in L15 clusters (Fig. 4A–G). The transcriptional repressor *Tramtrack* (*ttk*), known to inhibit EE cell fate^30,31^, was broadly expressed across many L15 clusters. Conversely, transcriptional markers associated with non-intestinal lineages were not enriched. These included markers of muscle development, MyosinD homolog *nautilus*^32^ and *twist*^33^, glial cells marker, *repo*^33,34^, glial cell master regulator *gcm*^35^ and the neural precursor gene *asense*^36^. Similar results were observed for L10, and L18. (Supplementary Fig. 3A-G and 4A-G). Altogether our analyses indicate that the clusters in the three cell lines were enriched for genes expressed in the intestine, with many of the clusters bearing the signatures of ISCs, suggesting all three cell lines were derived from the intestine.

**Figure 4.**
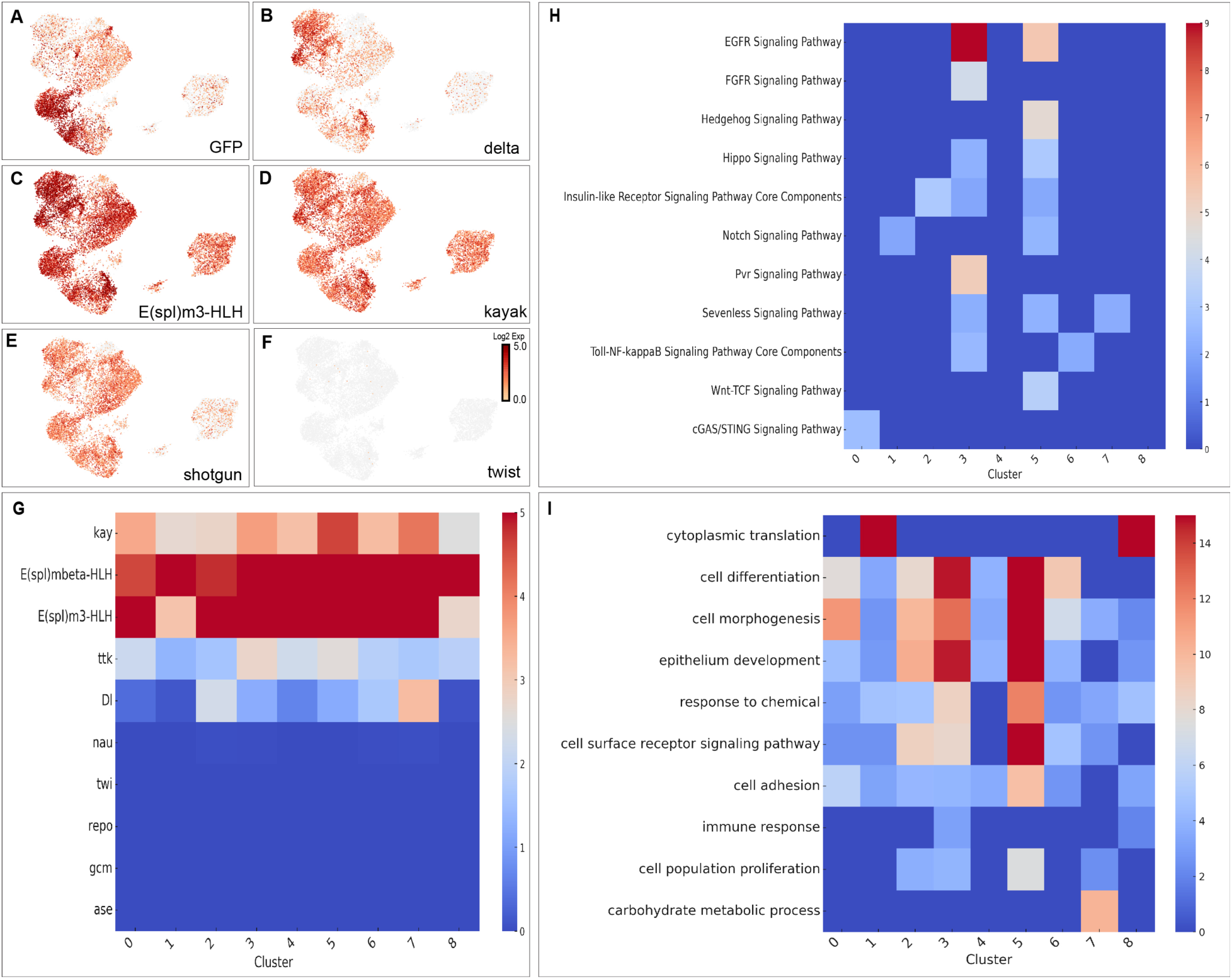
Gene Expression Analysis in L15. Gene expression feature plots for L15 single cell transcriptome analysis representing the expression of the following genes (A) *GFP*, (B) *delta*, (C) *E(spl)m3-HLH*, (D) *kayak*, (E) *shotgun*, and (F) *twist*. The relative expression level of each gene was indicated by red color, displayed as a natural logarithmic scale. (G) A table of gene expression plots compiled for L15 clusters. Cluster numbers are on the x-axis, and the gene names are on the y-axis. 2-color scale displays the gene’s median-normalized average (MNA) in each cluster. (H) Gene Set Enrichment Analysis (GSEA) Heatmap for signaling pathways enriched in L15 clusters. Cluster numbers are on the x-axis, and the identity of various signaling pathways are on the y-axis. (I) Gene Set Enrichment Analysis (GSEA) Heatmap for Biological Process Gene Ontology (GO) terms enriched in L15 clusters. Cluster numbers are on the x-axis, and the identity of various Biological Process GO terms are on the y-axis. The enrichment significance for both were displayed as negative log10 p-value scales.

## Multiple signaling pathways and distinct biological processes highlight the heterogeneity within the intestine cell lines

To investigate the extent of cell cluster-heterogeneity within a cell line, we utilized the gene set enrichment analysis using the PANGEA (Pathway, Network, and Gene-set Enrichment Analysis)^37^ platform to look at both signaling pathways and Gene Ontology (GO) term enrichment. In L15, clusters 3, EB like, and 6, ISC like, showed enrichment for genes involved in Toll-NFκB pathway and cluster 0, ISC like, was enriched for genes of the cGAS/STING pathway. Both Toll-NFκB and cGAS/STING pathways are involved in antiviral immunity in the *Drosophila* intestine^38^. Genes involved in Wnt-TCF signaling (cluster 5), Notch signaling (clusters 1 and 5), Hippo signaling (clusters 3 and 5), Hedgehog signaling (cluster 5) and Insulin signaling (clusters 2, 3 and 5) were also represented (Fig. 5H). Notably, clusters 3 and 5, both expressing markers for ISCs, were enriched for EGFR signaling, which is essential for the proliferation of larval AMPs and adult ISCs^39^. Likewise, in L10, the genes involved in the following signaling pathways including BMP (cluster 3, 6 and 13), JAK-STAT (cluster 1), Imd, and cGAS/STING pathways (cluster 10) were enriched, reflecting roles in gut epithelial development ^39,40^ and antiviral immune signaling ^38^ (Supplementary Fig. S3H). In L18, cluster 3, EE like, was enriched for genes in the Imd, Insulin, TNF-α, and Toll pathways, while cluster 9, ISC, showed enrichment for EGFR and Notch signaling components (Supplementary Fig. S4H). Genes in the Hedgehog pathway were enriched in cluster 5, EC like, while genes in the JAK-STAT pathway were enriched in cluster 6, ISC like. All together, the results suggest that various intestinal transcriptional profiles are active in these cell lines.

**Figure 5:**
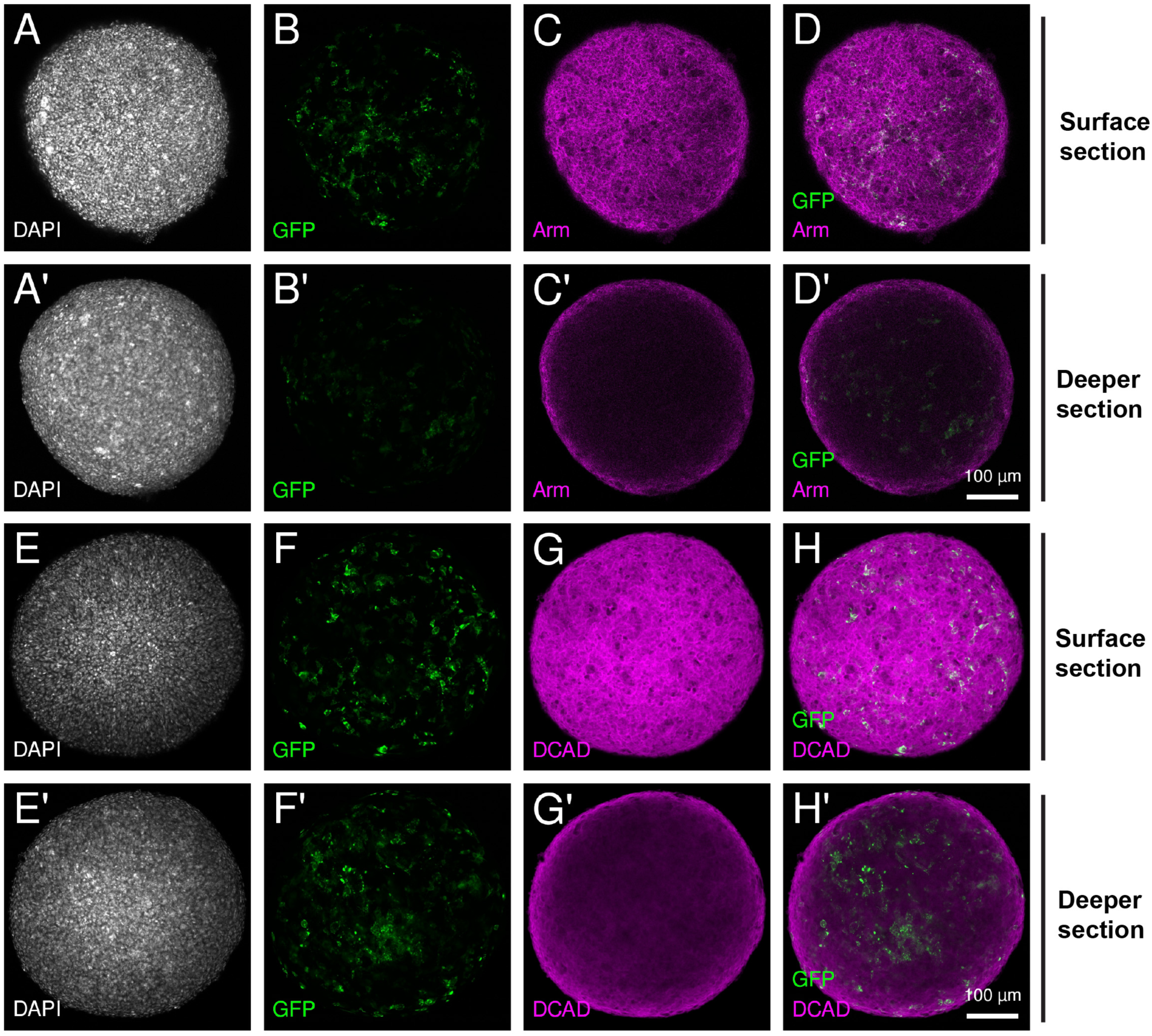
Cell adhesion molecules are expressed differentially within L15 three dimensional spheroids. 7-day hanging-drop culture derived L15 spheroids were fixed, immunostained and imaged. Either the surface (A-D, E-F) or deeper (A’-D’, E’-H)’ sections are shown for each spheroid. Immunostainings were performed for either Armadillo (Arm, C, C’) or DE-Cadherin (DCAD, G, G’). The speroids were also imaged for DAPI (A, A’, E, E’) and GFP (B, B’, F, F’). Merged images of either GFP and Arm (D, D’) or GFP and DCAD (H, H’) are also shown. Scale bar = 100 µm

We next investigated the Gene Ontology (GO) Biological Process represented in the clusters of the three cell lines. In L15, the majority of clusters were enriched in biological processes represented in intestinal functions^41^, including epithelium development (clusters 0-8), cell adhesion (clusters 0-8), cell morphogenesis (clusters 0-8), cell differentiation (clusters 0-6), response to chemicals (clusters 0-8), carbohydrate metabolic processes (cluster 7), and immune response (Fig. 4I). Some biological processes were only represented in specific clusters, for example clusters 1 and 8 were enriched in genes involved in cytoplasmic translation (Fig. 4I). In L10, similar analyses revealed an enrichment for genes involved in the antifungal immune response (cluster 10, EC, and 11, ISCs), and the Tachykinin signaling pathway (cluster 4 and 17, ISCs), reported to be involved in regulating lipid metabolism in the intestine^42^. Genes involved in endoderm formation cluster 6, ISCs, intestinal epithelial structure maintenance (Cluster 1, ISCs) and response to reactive oxygen species (cluster 8, ISCs) were enriched in L10 (Supplementary Fig. 3I). In L18, we found a significant enrichment for genes involved in the immune response in cluster 3 (EC cells), stem cell differentiation in clusters 4 (EB) and 7(differentiating EC), ribosome assembly in cluster 5 (EC) and cell cycle regulation in cluster 6 (ISCs) (Supplementary Fig. 4I). Taken together, the signaling pathway enrichment analysis demonstrated the heterogeneity of these lines and identified major signaling pathways relevant to intestinal development.

Complementarily, GO analysis revealed a complex scenario where distinct biological processes were isolated in specific clusters and occurring simultaneously and independently in each cell line.

We extended the gene enrichment analysis to include conducted GO Biological Process analysis on the differentially expressed markers genes within the clusters to identify biological processes associated with intestinal functions. In majority of the L15 clusters, we identified genes enriched in biological processes represented in intestinal functions^41^, including epithelium development (Clusters 0-8), cell adhesion (Clusters 0- 8), cell morphogenesis (Clusters 0-8), cell differentiation (Clusters 0-6), response to chemicals (Clusters 0-8), carbohydrate metabolic processes (Cluster 7), immune response (Fig. 4I). Some biological processes were highly represented in specific clusters, for example clusters 1 and 8 were enriched in genes involved in cytoplasmic translation (Fig. 4I). In L10, similar analyses revealed an enrichment for genes involved in the antifungal immune response (Cluster 10-11), Tachykinin signaling pathway (Cluster 4 and 17), reported to be involved in regulating lipid metabolism in the intestine^42^, endoderm formation (Cluster 6), and intestinal epithelial structure maintenance (Cluster 1) and response to reactive oxygen species (Cluster 8) (Supplementary Fig. 3I). In L18, we found a significant enrichment for genes involved in the immune response in in Cluster 3 (ISCs), stem cell differentiation in Clusters 4 (ISC) and 7(cardia), ribosome assembly is Cluster 5 (ISC) and cell cycle regulation in Cluster 6 (ISCs) (Supplementary Fig. 4I).

## L15 cells possess the ability to assemble into spheroids

As evident from the L15 scRNA seq analyses (Fig. 3 and Fig. 4), the L15 cell line is made up of several clusters with unique gene-expression profiles that possibly represent some of the physiologically relevant intestinal cell types. We therefore investigated the predisposition of this line to form spheroids and if these cell clusters can potentially organize into gut epithelium-like configuration^43^. When L15 cells were suspended in 20 ml hanging drops for seven days, they formed spheroids (Supplementary Fig. 5A-C). The spheroids were grown for up to 21 days with a media change every seven days (Supplementary Fig. 5D-E”). The spheroids were larger in volume at 21 days as compared to 7 days (Supplementary Fig. 5D, D”, E, E”), indicating growth within the spheroids over time. To this end, immunostaining the spheroids with an antibody against mitotic marker, phosphohistone H3 revealed mitotically active cells in multiple layers of the spheroids (Supplementary Fig. 6). These results demonstrate that the L15 cell line, when provided appropriate conditions, can assemble into mitotically active spheroids. This led us to investigate if any aspects of the adult intestinal epithelium are replicated in the spheroids.

The fly intestinal epithelium is a single layer of cells that possess intercellular adherens junctions (AJ) between EE cells^43^. To investigate if L15-derived spheroids form AJs, the localization of Armadillo (Arm) and DE-cadherin (DCAD) were characterized. Both Arm and DCAD are expressed only in the surface cell layer (Fig. 5A-D, E-H) of the spheroids and not in the deeper cells (Fig. 5A’-D’, E’-H’). However, GFP expression was observed in both surface cells (Fig. 5B, F) and deeper cells (Fig. 5B’, F’) in the same spheroid.

These results indicate formation of an epithelial-like surface exposed to the medium. The adult fly epithelium has a unique arrangement with the AJ more basal to septate junction (SJ)^43^. To explore this further, we investigated the localization of two other adult intestinal epithelial markers, Discs large (Dlg, a component of the SJ) and fly Talin ortholog, Rhea (an integrin adaptor and component of the basal layer)^43^. Both Talin (Supplementary Fig. 7A) and Dlg (Supplementary Fig. 7B) were localized to the surface exposed cells. Interestingly, while Dlg and DCAD (Supplementary Fig. 7B, B’) were found to be localized at similar confocal planes, Talin was localized more basally as compared to DCAD (Supplementary Fig. S7A, A’). These results with the 3D spheroids are in line with the organization of the cell junction markers in the adult midgut epithelium.

We next investigated if the distinct cellular layers in the L15-derived spheroids are associated with cell-type specific markers of intestinal epithelium. The spheroids were immunostained with Delta (Dl), Horse radish peroxidase (HRP), Nubbin (Nub), and Prospero (Pros). The ISC marker Dl is expressed in cells across the spheroids (Supplementary Fig. 8C). Cells positive for the ISC and EB marker, HRP, were interspersed throughout the spheroids (Supplementary Fig. 8G). The EC marker, Nub was only detected in very few cells per spheroid and these Nub positive cells were found deeper in the spheroid (Supplementary Fig. 8K). However, the EE marker Pros, was enriched predominantly in the surface-exposed cells (Supplementary Fig. 10).

From these results, it appears that expression of most of the intestinal cell-type specific markers in the L15-derived spheroid is not limited to distinct morphological structures within the spheroids. Moreover, these experiments demonstrate that given the appropriate conditions, L15 cells can assemble into complex entities that have distinct cellular layers with the surface exposed cells forming cellular junctions that mimic adult gut epithelium.

## Discussion

In this study, we report the creation of multiple *Drosophila melanogaster* embryonic intestine-derived cell lines leveraging the strategy of tissue-specific constitutively active Ras85D (Ras^V12^) expression^8^. As evident from the embryonic CG10116 GAL4 driven GFP expression and the single cell sequencing analyses, this approach has resulted in heterogenous cell lines that have a distinct intestinal gene expression signature. The protein product of CG10116 is predicted to be a lipase^44^, demonstrated to be expressed in the entire adult intestine^45^ and also shown to be expressed broadly in the embryonic intestine in this work. The putative function and the wildtype expression pattern of CG10116 therefore conforms with the observation that multiple clusters with distinct gene expression patterns exist in our embryonic intestine derived cell lines. This heterogeneity exemplifies the variety in physiological intestinal cell populations. For instance, despite being labelled as ISC, significant variance in the gene expression profiles of ISC’s from the different regions of the midgut has been characterized^46^. To evaluate the utility of the embryo-derived intestinal cell lines generated in this study, comparing the gene-expression profiles of these cell lines to the gut tissue across the various developmental stages would be relevant. However, these comparisons need to be viewed as indicative rather than absolute determinants for cells in culture.

In the developing *Drosophila* embryo, distinct cellular lineages that are precursors of the larval gut epithelium are specified by stage 16^47,48^. Moreover, endoderm-derived AMPs are also specified in the embryonic stage 11^49^. The embryo-derived larval midgut epithelium undergoes degeneration during pupal development^25^ and the AMPs begin to differentiate, to eventually form the adult midgut epithelium^50^. With this background, in the current study, we have utilized embryos from stage 16/17 to generate the intestinal cell lines. As such, we would expect a heterogenous population of cell lines that potentially are a combination of both either larval and adult precursors or differentiated populations. Single cell sequencing, followed by multiple downstream analyses of the generated cell lines revealed: a) expression of intestinal markers. For example, *kayak*, a key transcription factor and a spatial regulon in embryonic and larval midgut differentiation^29,51^. b) a combination of key biological pathways and processes involved in ISC specification, intestinal epithelial maintenance, and immune response were enriched in the derived cell lines. For example, we identified distinct clusters enriched for key pathways including Toll-NFκB, cGAS/STING, Wnt-TCF, Notch, Hippo, Insulin, and EGFR signaling pathways. The EGFR, Wnt and Notch pathways are essential for stem cell maintenance and cell fate specification, during intestinal epithelial renewal growth^25,52^. Hippo and insulin signaling have been shown to be required in controlling proliferation and the metabolic responses in the gut across multiple cell types ^53,54^.

These findings recapitulate the compartmentalized and cooperative nature of signaling pathways in shaping intestinal development. Overall, the three independent cell line, L15, L10 and L18 displayed relatively similar enrichment profiles for signaling pathways and biological processes.

The *Drosophila* larval/adult intestinal tissue is a complex three dimensional structure ^5^. We therefore investigated the tractability of the L15 cell line to form complex structures.

Upon providing constraints of a hanging drop, L15 cells were able to organize into 3D spheroids. The inherent cellular heterogeneity in the intestinal cell lines is most likely critical for spheroid formation. The medium exposed surface of the L15 spheroids selectively expressed both SJ and AJ markers. Moreover, the organization of the cell junction markers with the AJ localized more basal than SJ mimics the unique cell junction organization in the *Drosophila* adult midgut epithelium. This arrangement of the the cell junctions is not found in any other *Drosophila* epithelia ^43,55^. The finding that 3D spheroids have the similar cell junctional arrangement as the adult gut epithelium, offers the possibility that the L15 cell line can be utilized as a more complex tool than just cells in 2D culture. For instance, the effects of disrupting cell function using genetic and/or chemical approaches in the surface exposed cells on deeper cells can be investigated. These complex, heterogenous embryo-derived cell lines are potential resources for multiple areas of research.

Insects are one of the significant biotic causes for agricultural crop loss^56^, which is predicted to be exacerbated in the future^57^. However, only 9 of the 31 orders of insects are phytophagous, with almost all of the larvae from the order Lepidotera being phytophagous^58^. The need therefore to generate tools to understand basic biology of larval midgut epithelium and develop novel pest management strategies has led to the increased focus towards generating larval midgut cell lines, of which there are currently 13 cell lines listed in Cellosaurus^59^ and database for crop pest cell lines^60^. Of the 13 cell lines, 12 are derived from Lepidoteran larval midgut and the a single line is from Coleoptera^60^. These cell lines have been used to identify a novel insect infecting virus^61,62^, investigate effects of bacterial toxin^62,63^, recombinant protein production^64^, and response to prolonged hormone exposure^65^. The intestinal cell lines derived in this study can contribute to all of the above research questions/areas with the added advantage of possessing attP sites that can be utilized to generate transgenic cell lines with either orthologous or non-orthologous gene of interest from insect pests. Moreover, easy access to *Drosophila*-specific RNAi and gRNA libraries provide a platform to perform high-throughput genetic screens using these cell lines. Adult midgut-like characteristics observed upon manipuilation of the embryo-derived intestinal cell lines provide an important avenue to investigate processes that control development of the distinct larval and adult midgut epithelia. In summary, these lines fill the gap in the *Drosophila* toolkit, offering scalable and genetically manipulatable *in vitro* models to complement *in vivo* investigations of intestinal biology and toxicology.

## Methods

### Fly stocks, husbandry and embryo collection

The following fly stocks were used for the cross: [CG10116(-KDRT)GAL4]attP40; [20XUAS-6XGFP]attP2 (gift from Nick Sokol) and [UAS-Ras85D.V12], [att.w+.attP]JB89B (RRID:BDSC_64197). The resulting embryos had the genotype of [CG10116(-KDRT)GAL4]attP40 / +; [20XUAS-6XGFP]attP2/ UAS-Ras85D.V12], [att.w+.attP]JB89B (Figure 1A). CG10116-GAL4 was created by the removal of KD Recombinase Target (KDRT) sites flanking a stop codon cassette, by the KD recombinase encoded by gene A from *Kluyveromyces lactis*.

Flies were maintained on standard cornmeal-agar medium at 25°C. To collect embryos, chambers containing approximately 400 female and 200 male flies each were set up.

Embryos were collected for six hours and then aged for an additional 18 hours before processing. Thus, the collection was enriched for embryonic stages 16-17.

## Embryo and adult gut fixation and imaging

Embryos were collected and dechorionated in 50% bleach (∼3% hypochlorite concentration) for 3-4 minutes, and then rinsed in 1X PBS to remove bleach. Embryos were subsequently fixed on a nutator for 30 minutes in a glass scintillation vial containing 2 mL of. 1:1 fixative containing n-Heptane and 4% paraformaldehyde in 1XPBS. After fixation step, the lower formaldehyde layer (1 mL) was removed and 1 mL of methanol was added. The embyos in the heptane and methanol mixture was shaken, after which embryos will settle to the lower methanol phase. Emrbyos were then collected and strored in methanol at -20°C until ready for staining. Prior to staining, embryo samples were rehydrydated in a mixture of methanol: 1X PBS mix (3:1, 1:1, and 1:3) progressively. Each incubation period was for 15 minutes.

Adult guts were dissected in 1X PBS and immediately fixed for 15 minutes in 4% paraformaldehyde solution containing 1X PBS. Imaging for emrbyos and adult guts follow similar procedures. Briefly, samples were mounted using Vectashield H-1200 mounting media containing DAPI and imaged using Zeiss Axio Observer microscope with 10X objective.

## Embryo processing and homogenization

Embryos were rinsed with deionized filtered water using a squirt bottle and collected in a 40-μm sieve to remove large clumps and debris. The embryos were transferred to a 15 mL conical tube containing approximately 5 mL of TXN (0.7% NaCl and 0.02% Triton X-100) and washed multiple times with deionized filtered water to eliminate extraneous material. After washing, embryos were incubated in a solution of 50% bleach for 5 minutes to remove the chorion. Following chorion removal, embryos were washed several times with TXN to ensure complete removal of bleach. Embryos were then transferred to a glass homogenizer with TXN and homogenized with 2-3 strokes in 0.2% Trypsin (Thermo Fisher 12563011). Homogenate was transferred to a 15 mL conical tube and allowed to settle at room temperature for 30-35 minutes. The homogenate was centrifuged at 700 x g for 10 minutes, and the supernatant was discarded. The cell pellet was resuspended in 1-3 mL of MB10+Antibiotics: M3 (Sigma S8398) supplemented with 0.25% bactopeptone (Difco 211677), 0.1% yeast extract (Sigma Y1000), 10% Fetal Bovine Serum FBS (Hyclone Cytiva SH30070.03), and 1:100 of antibiotic and antimycotic mixture (Sigma A5955). The suspension was passed through a 50-micron filter to remove debris and large particles and was seeded into a 12.5 cm^2^ cell culture (T-12.5) flask at a final volume of 3 mL.

## Primary cell cultures and maintenance

Primary cultures were maintained in the same T-12.5 flask without disturbing them for 3- 5 days to allow cells to attach and proliferate. Cultures were periodically checked for contaminants. The media was replenished or refreshed every two weeks with 0.75 to 1 ml of MB10+Antibiotics. The primary cultures reached 90% confluency and contained GFP positive cluster of cells in 4-5 weeks. At this point, the cultures were passaged 1:2 into additional T12-5 flasks and subsequently expanded.

## Culture conditions for new continuous cell lines

Cells were grown in 25 cm^2^ T-flasks at 25°C in M3 + 10% FBS (M10) medium and were passaged at between 90% and full confluence using trypsin to release cells from the tissue-culture surface. Trypsin was essential since all the lines were adherent. In cases where trypsin treatment did not dislodge the monolayer after longer than 10 minutes, cell scrapers were used to aid dissociation. Cells were pelleted and sub-cultured at a ratio between 1:4 to 1:5. Cells were checked using an inverted microscope approximately every 2-3 days. Cells were passaged every 5–7 days based on growth rates.

## Freeze/thaw protocol

Cells from healthy confluent cultures were dislodged from the growth surfaces, pelleted and resuspended in 0.5 mL of the freezing media made of M3 medium supplemented with 20% heat inactivated fetal bovine serum and 10% DMSO (Dimethyl sulfoxide).

Cryopreservation was done at multiple time points to ensure a steady supply of cell lines with similar passage numbers were used in experiments. L10 was cryopreserved in bulk at passages 11, 12, 17. L15 was cryopreserved in bulk at passages 13, 16, and 47. L18 was cryopreserved in bulk at passages 14, 19 and 29. Early passages of these lines (between passages 2-8) were also available in lower quantities.

As with all tissue culture systems, maintaining frozen stocks of cell lines at low passage numbers is essential to preserve their original characteristics^3^. To support reproducibility and consistency, we have made batches of aliquots of the three cell lines used in our scRNAseq analyses at early passage numbers comparable to those used for subsequent analysis.

## Growth curve analysis

Cell cultures were grown in M10. To analyze growth, cultures were seeded at a starting cell count of 5x10^5^ cells/ml (L10 and L15) or 1x10^5^ cells/ml (L18) and distributed into wells of a 24-well plate. Aliquots were drawn from the wells at between 24-50 hour intervals for ∼200 hours and cell count measured. Growth data from four biological replicates for each intestinal line was collected and compiled. Doubling times were calculated using https://www.doubling-time.com/compute.php.

### Flow Cytometry

2x10^6^ cells were collected from log phase cultures and fixed in 70% EtOH and stored at 4°C. On the day of analysis, the fixed cells were washed with ice cold 1x PBS and stained with Propidium Iodide solution (1x PBS, 0.1% Tirion-X 100, 20mg/ml propidium iodide, 0.2mg/ml Rnase A) for 15 minutes at 37°C. Samples were processed using either a BD FACSAria IIu (L15) or a CytoFLEX LX (L10 and L18) with a laser excitation of 610-nm at the Indiana University Bloomington Flow Cytometry Core Facility. At least 30,000 events were recorded for 3 biological replicates for each intestinal line. Samples were analyzed using floreada.io.

## Single-cell 3’ RNA-seq

Single-cell 3’ gene expression assay was conducted using the Chromium single cell system (10x Genomics, Inc) and NovaSeq 6000 sequencer (Illumina, Inc). Every cell suspension was first checked for cell number, cell viability. Depending on the quality of the initial cell suspension, the single cell preparation included re-suspension, centrifugation, and filtration to remove cell debris, dead cells and cell aggregates. The final single cell suspension (>2 million cells/mL) had high viability (>88%) and minimal cell debris and aggregation. Appropriate number of cells (L10: 16503 cells, L15: 17139 cells, L18: 19564 cells) were loaded on a multiple-channel micro-fluidics chip of the Chromium Single Cell Instrument (10x Genomics). Single cell gel beads in emulsion containing barcoded oligonucleotides and reverse transcriptase reagents were generated with the Next Gen single cell reagent kit (10X Genomics). Following cell capture and cell lysis, cDNA was synthesized and amplified. At each step, the quality of cDNA and library was examined by Bioanalyzer. The resulting dual-indexed library was sequenced on Illumina NovaSeq 6000. 28-bp reads including cell barcode and UMI sequences and 100-bp RNA reads were generated. The data discussed in this publication have been deposited in NCBI’s Gene Expression Omnibus ^66^ and are accessible through GEO Series accession number GSE295099 (https://www.ncbi.nlm.nih.gov/geo/query/acc.cgi?acc=GSE295099). The reviewer token for private access during the manuscript review will be made available for the journal editor.

## Single-cell data analysis

Single-cell RNA-seq data was processed using CellRanger 7.0.1 (http://support.10xgenomics.com/). The feature-cell barcode matrices generated from CellRanger were used for further analysis with the R package Seurat v4^67–69^. After quality check and normalization, the single-cell data from multiple samples was integrated for dimensionality reduction, graph-based cell clustering analysis and cluster marker gene identification. To annotate each cell cluster, we combined both manual interpretation with known canonical marker genes, cluster marker genes, and annotation with publicly available fly cell atlas single cell dataset (whole body_dm_FCAvs2_10x and Intestine_dm_FCAvs2_10x)^28^.

## Three-dimensional cell culture

Confluent cultures of the intestinal cell lines were dispersed, and the cell count estimated. 20 µl of the cultures at 0.5 x10^6^ cells/ml were aliquoted into the reverse side of 35 mm culture dish lids. Seven 20 µl drops were aliquoted onto each lid. The lids were carefully turned over onto the bottom of the dish containing 1 ml of media. The cultures were incubated for either 7, 14 or 21 days with a weekly media change for cultures longer than 7 days. To acquire brightfield images, the lids were carefully turned over and imaged with a Nikon TS1 microscope.

## Immunofluoresence

To fix spheroids, an equal volume (20 µl) of 8% paraformaldehyde (PF) in 2X Phosphate Buffered Saline (PBS) was added to the cultures (for a final 4% PF concentration in 1X PBS) for 20 minutes. The spheroids in the PF fix solution were transferred to a 0.5 ml tube and the fix removed. The fixed spheroids were washed twice with 0.1% Triton X-100 in phosphate-buffered saline (PBT). Subsequently the fixed spheroids were incubated in PBT for 10 min at room temperature before immunostaining. To immunostain the 3D spheroids, they were first blocked with Blocking solution (10% Normal Goat Serum (Abcam, AB7481), 1% BSA, 0.1% Triton in PBS). The spheroids were then incubated with primary antibodies in Blocking solution overnight at 4°C, followed by five washes in PBT. After completion of the washes, the spheroids were incubated in the appropriate secondary antibodies diluted in 3D- Blocking solution for one hour at room temperature. The spheroids were then subjected to five washes in PBT, following which they were mounted in Prolong Glass Antifade Mountant (Invitrogen, P36984), allowed to cure overnight at room temperature before confocal imaging in the Leica SP8 microscope at Light Microscopy Imaging Center at Indiana University, Biology Department. Confocal images were analyzed and the final figures prepared with either Open Microscopy Environment webserver (https://www.openmicroscopy.org/index.html,^70^), Fiji^71^ or Adobe Illustrator.

The following primary antibodies were used to immunostaining: rat anti-DCAD2 (DSHB Cat# DCAD2, RRID:AB_528120)^72^, mouse anti-Armadillo (DSHB Cat# N2 7A1 ARMADILLO, RRID:AB_528089)^73^, mouse anti-Prospero (DSHB Cat# Prospero, RRID:AB_528440)^74^, mouse anti-Delta (DSHB Cat# c594.9b, RRID:AB_528194)^75^, mouse anti-Nubbin (DSHB Cat# Nub 2D4, RRID:AB_2722119)^76^, mouse anti-discs large (DSHB Cat# 4F3 anti-discs large, RRID:AB_528203)^77^, anti-talin (DSHB Cat# Talin A22A, RRID:AB_10660289) ^78^(all DSHB antibodies were used at 2 µg/ml), goat anti-Horseradish Peroxidase-Alexa647 (Jackson ImmunoResearch Cat# 123-605-021, RRID: AB_2338967, 1:1000), and rabbit anti-phospho-histone H3 (Cell Signaling Technology, Cat# 9701S). For immunostaining spheroids, the following secondary antibodies were used at a dilution of 1:250 in Blocking solution: Alexa647-conjugated goat anti-mouse (Jackson ImmunoResearch Cat# 115-605-166, RRID: AB_2338914), Alexa594-conjugated goat anti-rat (Jackson ImmunoResearch, Cat# 112-585-003, RRID: AB_2338372), Alexa594-conjugated goat anti-mouse (Jackson ImmunoResearch Cat# 115-585-166, RRID: AB_2338883), or Alexa633-conjugated goat anti-rabbit (Thermo Cat# A-21070, RRID: AB_2525731). Secondary antibody incubations also contained DAPI at 1:5000 from 5 mg/ml stock.

## Acknowledgements

Work at the Drosophila Genomics Resource Center is supported by NIH grant P40OD010949. Microscopy work was carried out at the Light Microscopy Imaging Center (LMIC) at Indiana University Bloomington. Sequencing analysis was carried out at the Center for Medical Genomics at Indiana University School of Medicine, which is partially supported by the Indiana University Grand Challenges Precision Health

Initiative. Flow cytometry was done at IUB Flow Cytometry Core Facility. We used the FACSAria IIu and the CytoFLEX LX instrument (This equipment was funded in part by the IU Office of Research through the Research Equipment Fund.) Stocks obtained from the Bloomington Drosophila Stock Center (NIH P40OD018537) were used in this study. We used FlyBase (release FB2025_02) to find information on function, stocks and gene expression information. Fig. 2B was created with BioRender.com.

## Author contributions

AL, DM and AZ designed the study. AL and DM equally contributed as co-first authors. AL and DM supervised, performed experiments and data analysis. AL coordinated the processing of single cell transcriptomic samples and performed related analyses. DM developed 3D-culture conditions and imaging techniques related to spheroids. PB characterized ploidy and growth curves. LM and ME were involved in establishing the cell lines. AL, DM and AZ wrote the manuscript. All authors gave their permission for manuscript submission.

## Data availability statement

Cell lines generated during this study are available from the Drosophila Genomics Resource Center for public distribution. The datasets generated during and analysed during the current study are available in NCBI Gene Expression Omnibus and are accessible through GEO Series accession number GSE295099 (https://www.ncbi.nlm.nih.gov/geo/query/acc.cgi?acc=GSE295099).

## Competing interests

The authors declare no competing interests.

## Notes

### Competing Interest Statement

The authors have declared no competing interest.

## REFERENCES

1 Yee, C. M., Zak, A. J., Hill, B. D. & Wen, F. The Coming Age of Insect Cells for Manufacturing and Development of Protein Therapeutics. Ind Eng Chem Res 57, 10061–10070 (2018). 10.1021/acs.iecr.8b00985

2 He, X. et al. Insect Cell-Based Models: Cell Line Establishment and Application in Insecticide Screening and Toxicology Research. Insects 14 (2023). 10.3390/insects14020104

3 Luhur, A., Klueg, K. M. & Zelhof, A. C. Generating and working with Drosophila cell cultures: Current challenges and opportunities. Wiley Interdiscip Rev Dev Biol 8, e339 (2019). 10.1002/wdev.339

4 Capo, F., Wilson, A. & Di Cara, F. The Intestine of Drosophila melanogaster: An Emerging Versatile Model System to Study Intestinal Epithelial Homeostasis and Host-Microbial Interactions in Humans. Microorganisms 7 (2019). 10.3390/microorganisms7090336

5 Miguel-Aliaga, I., Jasper, H. & Lemaitre, B. Anatomy and Physiology of the Digestive Tract of Drosophila melanogaster. Genetics 210, 357–396 (2018). 10.1534/genetics.118.300224

6 Cherbas, L. et al. The transcriptional diversity of 25 Drosophila cell lines. Genome Res 21, 301–314 (2011). 10.1101/gr.112961.110

7 Stoiber, M., Celniker, S., Cherbas, L., Brown, B. & Cherbas, P. Diverse Hormone Response Networks in 41 Independent Drosophila Cell Lines. *G3* (*Bethesda*) 6, 683-694 (2016). 10.1534/g3.115.023366

8 Coleman-Gosser, N. et al. Continuous muscle, glial, epithelial, neuronal, and hemocyte cell lines for Drosophila research. Elife 12 (2023). 10.7554/eLife.85814

9 Bravo, A., Gill, S. S. & Soberon, M. Mode of action of Bacillus thuringiensis Cry and Cyt toxins and their potential for insect control. Toxicon 49, 423–435 (2007). 10.1016/j.toxicon.2006.11.022

10 ffrench-Constant, R. H., Dowling, A. & Waterfield, N. R. Insecticidal toxins from Photorhabdus bacteria and their potential use in agriculture. Toxicon 49, 436–451 (2007). 10.1016/j.toxicon.2006.11.019

11 Gruber, C. W., Cemazar, M., Anderson, M. A. & Craik, D. J. Insecticidal plant cyclotides and related cystine knot toxins. Toxicon 49, 561–575 (2007). 10.1016/j.toxicon.2006.11.018

12 King, G. F. Modulation of insect Ca(v) channels by peptidic spider toxins. Toxicon 49, 513–530 (2007). 10.1016/j.toxicon.2006.11.012

13 Gordon, D. et al. The differential preference of scorpion alpha-toxins for insect or mammalian sodium channels: implications for improved insect control. Toxicon 49, 452–472 (2007). 10.1016/j.toxicon.2006.11.016

14 Wu, C., Chakrabarty, S., Jin, M., Liu, K. & Xiao, Y. Insect ATP-Binding Cassette (ABC) Transporters: Roles in Xenobiotic Detoxification and Bt Insecticidal Activity. Int J Mol Sci 20 (2019). 10.3390/ijms20112829

15 Rosner, J., Tietmeyer, J. & Merzendorfer, H. Functional analysis of ABCG and ABCH transporters from the red flour beetle, Tribolium castaneum. Pest Manag Sci 77, 2955–2963 (2021). 10.1002/ps.6332

16 Justiniano, S. E., Mathew, A., Mitra, S., Manivannan, S. N. & Simcox, A. Loss of the tumor suppressor Pten promotes proliferation of Drosophila melanogaster cells in vitro and gives rise to continuous cell lines. PLoS One 7, e31417 (2012). 10.1371/journal.pone.0031417

17 Simcox, A. et al. Efficient genetic method for establishing Drosophila cell lines unlocks the potential to create lines of specific genotypes. PLoS Genet 4, e1000142 (2008). 10.1371/journal.pgen.1000142

18 Buddika, K., Xu, J., Ariyapala, I. S. & Sokol, N. S. I-KCKT allows dissection-free RNA profiling of adult Drosophila intestinal progenitor cells. Development 148 (2021). 10.1242/dev.196568

19 Hung, R. J. et al. A cell atlas of the adult Drosophila midgut. Proc Natl Acad Sci U S A 117, 1514–1523 (2020). 10.1073/pnas.1916820117

20 Bateman, J. R., Lee, A. M. & Wu, C. T. Site-specific transformation of Drosophila via phiC31 integrase-mediated cassette exchange. Genetics 173, 769–777 (2006). 10.1534/genetics.106.056945

21 Manivannan, S. N. et al. Targeted Integration of Single-Copy Transgenes in Drosophila melanogaster Tissue-Culture Cells Using Recombination-Mediated Cassette Exchange. Genetics 201, 1319–1328 (2015). 10.1534/genetics.115.181230

22 Takashima, S. et al. Development of the Drosophila entero-endocrine lineage and its specification by the Notch signaling pathway. Dev Biol 353, 161–172 (2011). 10.1016/j.ydbio.2011.01.039

23 Chen, J. & St Johnston, D. Epithelial Cell Polarity During Drosophila Midgut Development. Front Cell Dev Biol 10, 886773 (2022). 10.3389/fcell.2022.886773

24 Ohlstein, B. & Spradling, A. The adult Drosophila posterior midgut is maintained by pluripotent stem cells. Nature 439, 470–474 (2006). 10.1038/nature04333

25 Jiang, H. & Edgar, B. A. EGFR signaling regulates the proliferation of Drosophila adult midgut progenitors. Development 136, 483–493 (2009). 10.1242/dev.026955

26 Korzelius, J. et al. Escargot maintains stemness and suppresses differentiation in Drosophila intestinal stem cells. EMBO J 33, 2967–2982 (2014). 10.15252/embj.201489072

27 Hu, Y. et al. DRscDB: A single-cell RNA-seq resource for data mining and data comparison across species. Comput Struct Biotechnol J 19, 2018–2026 (2021). 10.1016/j.csbj.2021.04.021

28 Li, H. et al. Fly Cell Atlas: A single-nucleus transcriptomic atlas of the adult fruit fly. Science 375, eabk2432 (2022). 10.1126/science.abk2432

29 Souid, S. & Yanicostas, C. Differential expression of the two Drosophila fos/kayak transcripts during oogenesis and embryogenesis. Dev Dyn 227, 150–154 (2003). 10.1002/dvdy.10293

30 Guo, X., Wang, C., Zhang, Y., Wei, R. & Xi, R. Cell-fate conversion of intestinal cells in adult Drosophila midgut by depleting a single transcription factor. Nat Commun 15, 2656 (2024). 10.1038/s41467-024-46956-8

31 Wang, C., Guo, X., Dou, K., Chen, H. & Xi, R. Ttk69 acts as a master repressor of enteroendocrine cell specification in Drosophila intestinal stem cell lineages. Development 142, 3321–3331 (2015). 10.1242/dev.123208

32 Misquitta, L. & Paterson, B. M. Targeted disruption of gene function in Drosophila by RNA interference (RNA-i): a role for nautilus in embryonic somatic muscle formation. Proc Natl Acad Sci U S A 96, 1451–1456 (1999). 10.1073/pnas.96.4.1451

33 Baylies, M. K. & Bate, M. twist: a myogenic switch in Drosophila. Science 272, 1481–1484 (1996). 10.1126/science.272.5267.1481

34 Xiong, W. C., Okano, H., Patel, N. H., Blendy, J. A. & Montell, C. repo encodes a glial- specific homeo domain protein required in the Drosophila nervous system. Genes Dev 8, 981–994 (1994). 10.1101/gad.8.8.981

35 Mao, H., Lv, Z. & Ho, M. S. Gcm proteins function in the developing nervous system. Dev Biol 370, 63–70 (2012). 10.1016/j.ydbio.2012.07.018

36 Brand, M., Jarman, A. P., Jan, L. Y. & Jan, Y. N. asense is a Drosophila neural precursor gene and is capable of initiating sense organ formation. Development 119, 1–17 (1993). 10.1242/dev.119.Supplement.1

37 Hu, Y. et al. PANGEA: a new gene set enrichment tool for Drosophila and common research organisms. Nucleic Acids Res 51, W419–W426 (2023). 10.1093/nar/gkad331

38 Nigg, J. C. et al. Viral infection disrupts intestinal homeostasis via Sting-dependent NF-kappaB signaling in Drosophila. Curr Biol 34, 2785–2800 e2787 (2024). 10.1016/j.cub.2024.05.009

39 Xu, N. et al. EGFR, Wingless and JAK/STAT signaling cooperatively maintain Drosophila intestinal stem cells. Dev Biol 354, 31–43 (2011). 10.1016/j.ydbio.2011.03.018

40 Tian, A. & Jiang, J. Intestinal epithelium-derived BMP controls stem cell self-renewal in Drosophila adult midgut. Elife 3, e01857 (2014). 10.7554/eLife.01857

41 Marianes, A. & Spradling, A. C. Physiological and stem cell compartmentalization within the Drosophila midgut. Elife 2, e00886 (2013). 10.7554/eLife.00886

42 Song, W., Veenstra, J. A. & Perrimon, N. Control of lipid metabolism by tachykinin in Drosophila. Cell Rep 9, 40–47 (2014). 10.1016/j.celrep.2014.08.060

43 Chen, J., Sayadian, A. C., Lowe, N., Lovegrove, H. E. & St Johnston, D. An alternative mode of epithelial polarity in the Drosophila midgut. PLoS Biol 16, e3000041 (2018). 10.1371/journal.pbio.3000041

44 Horne, I., Haritos, V. S. & Oakeshott, J. G. Comparative and functional genomics of lipases in holometabolous insects. Insect Biochem Mol Biol 39, 547–567 (2009). 10.1016/j.ibmb.2009.06.002

45 Ariyapala, I. S. et al. Identification of Split-GAL4 Drivers and Enhancers That Allow Regional Cell Type Manipulations of the Drosophila melanogaster Intestine. Genetics 216, 891–903 (2020). 10.1534/genetics.120.303625

46 Dutta, D. et al. Regional Cell-Specific Transcriptome Mapping Reveals Regulatory Complexity in the Adult Drosophila Midgut. Cell Rep 12, 346–358 (2015). 10.1016/j.celrep.2015.06.009

47 Hoppler, S. & Bienz, M. Specification of a single cell type by a Drosophila homeotic gene. Cell 76, 689–702 (1994). 10.1016/0092-8674(94)90508-8

48 Szuts, D., Eresh, S. & Bienz, M. Functional intertwining of Dpp and EGFR signaling during Drosophila endoderm induction. Genes Dev 12, 2022–2035 (1998). 10.1101/gad.12.13.2022

49 Hartenstein, A. Y., Rugendorff, A., Tepass, U. & Hartenstein, V. The function of the neurogenic genes during epithelial development in the Drosophila embryo. Development 116, 1203–1220 (1992). 10.1242/dev.116.4.1203

50 Takashima, S., Younossi-Hartenstein, A., Ortiz, P. A. & Hartenstein, V. A novel tissue in an established model system: the Drosophila pupal midgut. Dev Genes Evol 221, 69–81 (2011). 10.1007/s00427-011-0360-x

51 Wang, M. et al. High-resolution 3D spatiotemporal transcriptomic maps of developing Drosophila embryos and larvae. Dev Cell 57, 1271–1283 e1274 (2022). 10.1016/j.devcel.2022.04.006

52 Sun, H. et al. Wnt/beta-catenin signaling within multiple cell types dependent upon kramer regulates Drosophila intestinal stem cell proliferation. iScience 27, 110113 (2024). 10.1016/j.isci.2024.110113

53 Poernbacher, I., Baumgartner, R., Marada, S. K., Edwards, K. & Stocker, H. Drosophila Pez acts in Hippo signaling to restrict intestinal stem cell proliferation. Curr Biol 22, 389–396 (2012). 10.1016/j.cub.2012.01.019

54 Choi, N. H., Lucchetta, E. & Ohlstein, B. Nonautonomous regulation of Drosophila midgut stem cell proliferation by the insulin-signaling pathway. Proc Natl Acad Sci U S A 108, 18702–18707 (2011). 10.1073/pnas.1109348108

55 Tepass, U., Tanentzapf, G., Ward, R. & Fehon, R. Epithelial cell polarity and cell junctions in Drosophila. Annu Rev Genet 35, 747–784 (2001). 10.1146/annurev.genet.35.102401.091415

56 Oerke, E. C. Crop losses to pests. J Agr Sci-Cambridge 144, 31–43 (2006). 10.1017/S0021859605005708

57 Deutsch, C. A. et al. Increase in crop losses to insect pests in a warming climate. Science 361, 916–919 (2018). 10.1126/science.aat3466

58 Douglas, A. E. Strategies for Enhanced Crop Resistance to Insect Pests. Annu Rev Plant Biol 69, 637–660 (2018). 10.1146/annurev-arplant-042817-040248

59 Bairoch, A. The Cellosaurus, a Cell-Line Knowledge Resource. J Biomol Tech 29, 25–38 (2018). 10.7171/jbt.18-2902-002

60 Arya, S. K., Goodman, C. L., Stanley, D. & Palli, S. R. A database of crop pest cell lines. In Vitro Cell Dev Biol Anim 58, 719–757 (2022). 10.1007/s11626-022-00710-w

61 Pringle, F. M., Johnson, K. N., Goodman, C. L., McIntosh, A. H. & Ball, L. A. Providence virus: a new member of the Tetraviridae that infects cultured insect cells. Virology 306, 359–370 (2003). 10.1016/s0042-6822(02)00052-1

62 Li, J. et al. Establishment and characterization of a novel cell line from midgut tissue of Helicoverpa armigera (Lepidoptera: Noctuidae). In Vitro Cell Dev Biol Anim 51, 562–571 (2015). 10.1007/s11626-015-9870-6

63 Zhou, K. et al. Establishment of two midgut cell lines from the fall armyworm, Spodoptera frugiperda (Lepidoptera: Noctuidae). In Vitro Cell Dev Biol Anim 56, 10–14 (2020). 10.1007/s11626-019-00420-w

64 Davis, T. R. et al. Comparative recombinant protein production of eight insect cell lines. In Vitro Cell Dev Biol Anim **29A**, 388–390 (1993). 10.1007/BF02633986

65 Hu, W. et al. Morphological and molecular effects of 20-hydroxyecdysone and its agonist tebufenozide on CF-203, a midgut-derived cell line from the spruce budworm, Choristoneura fumiferana. Arch Insect Biochem Physiol 55, 68–78 (2004). 10.1002/arch.10124

66 Edgar, R., Domrachev, M. & Lash, A. E. Gene Expression Omnibus: NCBI gene expression and hybridization array data repository. Nucleic Acids Res 30, 207–210 (2002). 10.1093/nar/30.1.207

67 Stuart, T. et al. Comprehensive Integration of Single-Cell Data. Cell 177, 1888–1902 e1821 (2019). 10.1016/j.cell.2019.05.031

68 Hao, Y. et al. Integrated analysis of multimodal single-cell data. Cell 184, 3573–3587 e3529 (2021). 10.1016/j.cell.2021.04.048

69 Butler, A., Hoffman, P., Smibert, P., Papalexi, E. & Satija, R. Integrating single-cell transcriptomic data across different conditions, technologies, and species. Nat Biotechnol 36, 411–420 (2018). 10.1038/nbt.4096

70 Allan, C. et al. OMERO: flexible, model-driven data management for experimental biology. Nat Methods 9, 245–253 (2012). 10.1038/nmeth.1896

71. Schindelin, J., et al. Fiji: an open-source platform for biological-image analysis. Nat Methods 9, 676-682 (2012). 10.1038/nmeth.2019

72 Oda, H., Uemura, T., Harada, Y., Iwai, Y. & Takeichi, M. A Drosophila homolog of cadherin associated with armadillo and essential for embryonic cell-cell adhesion. Dev Biol 165, 716–726 (1994). 10.1006/dbio.1994.1287

73 Riggleman, B., Schedl, P. & Wieschaus, E. Spatial expression of the Drosophila segment polarity gene armadillo is posttranscriptionally regulated by wingless. Cell 63, 549–560 (1990). 10.1016/0092-8674(90)90451-j

74 Spana, E. P. & Doe, C. Q. The prospero transcription factor is asymmetrically localized to the cell cortex during neuroblast mitosis in Drosophila. Development 121, 3187–3195 (1995). 10.1242/dev.121.10.3187

75 Sun, X. & Artavanis-Tsakonas, S. The intracellular deletions of Delta and Serrate define dominant negative forms of the Drosophila Notch ligands. Development 122, 2465–2474 (1996). 10.1242/dev.122.8.2465

76 Averof, M. & Cohen, S. M. Evolutionary origin of insect wings from ancestral gills. Nature 385, 627–630 (1997). 10.1038/385627a0

77 Parnas, D., Haghighi, A. P., Fetter, R. D., Kim, S. W. & Goodman, C. S. Regulation of postsynaptic structure and protein localization by the Rho-type guanine nucleotide exchange factor dPix. Neuron 32, 415–424 (2001). 10.1016/s0896-6273(01)00485-8

78 Brown, N. H. et al. Talin is essential for integrin function in Drosophila. Dev Cell 3, 569–579 (2002). 10.1016/s1534-5807(02)00290-3

