## Supplementary File for "Establishment of *Drosophila* Intestinal Cell Lines"

### Supplementary Figure Legends

**Supplementary Figure S1. CG10116-GAL4 expression profile in the developing embryo.** Stages 11–17 embryos showing GFP expression (green) in the developing midgut. Each row displays a representative embryonic stage, indicated in the top left corner of the GFP column. The three columns show: (1) GFP fluorescence, (2) DAPI staining, and (3) merged channels. D/V/L labels in the merged column indicate the orientation of the embryo: dorsal (D), ventral (V), or lateral (L), shown in the top right corner of each image. A red arrowhead marks the GFP signal in the anterior midgut. Scale bar = 100 μm.

**Supplementary Figure S2. Morphological and Growth profiles of L10, L15 and L18**. (A) Brightfield micrograph of L10, (A') GFP and (A'') merged channels. (B) Brightfield micrograph of L15, (B') GFP and (B'') merged channels. (C) Brightfield micrograph of L18, (C') GFP and (C'') merged channels. Scale bar = 20 μm (D-F) Growth curves of L10, L15 and L18. (G-I) Cell ploidy of L10, L15, and L18. J. Table showing the doubling time of L10, L15, and L18.

**Supplementary Figure S3. Gene Expression Analysis in L10.** Gene expression feature plots for L10 single cell transcriptome analysis representing the expression of the following genes (A) *GFP*, (B) *delta*, (C) *E(spl)m3-HLH*, (D) *kayak*, (E) *shotgun*, and (F) *twist*. The relative expression level of each gene was indicated by red color, displayed as a natural logarithmic scale. (G) Table of gene expression plots compiled for L10 clusters. Cluster numbers are on the x-axis, and the gene names are on the y-axis. 2-color scale displays the gene's median-normalized average (MNA) in each cluster, i.e. averaged barcode size-normalized gene count. (H) Gene Set Enrichment Analysis (GSEA) Heatmap for signaling pathways enriched in L10 clusters. Cluster numbers are on the x-axis, and the identity of various signaling pathways are on the y-axis. (I) Gene Set Enrichment Analysis (GSEA) Heatmap for Biological Process Gene Ontology (GO) terms enriched in L10 clusters. Cluster numbers are on the x-axis, and the identity of various Biological Process GO terms are on the y-axis. The enrichment significance for both were displayed as negative log10 p-value scales.

**Supplementary Figure S4. Gene Expression Analysis in L18.** Gene expression feature plots for L18 single cell transcriptome analysis representing the expression of the following genes (A) *GFP*, (B) *delta*, (C) *E(spl)m3-HLH*, (D) *kayak*, (E) *shotgun*, and (F) *twist*. The relative expression level of each gene was indicated by red color, displayed as a natural logarithmic scale. (G) Table of gene expression plots compiled for L18 clusters. Cluster numbers are on the x-axis, and the gene names are on the y-axis. 2-color scale displays the gene's median-normalized average (MNA) in each cluster, i.e. averaged barcode size-normalized gene count. (H) Gene Set Enrichment Analysis (GSEA) Heatmap for signaling pathways enriched in L18 clusters. Cluster numbers are on the x-axis, and the identity of various signaling pathways are on the y-axis. (I) Gene Set Enrichment Analysis (GSEA) Heatmap for Biological Process Gene Ontology (GO) terms enriched in L18 clusters. Cluster numbers are on the x-axis, and the identity of various Biological Process GO terms are on the y-axis. The enrichment significance for both were displayed as negative log10 p-value scales.

**Supplementary Figure S5**. **The 3D organization of embryonic intestinal line L15.** Different cell counts; 0.175 x $10^6$ (A), 0.35 x $10^6$ (B), or 0.7 x $10^6$ (C) were incubated in hanging drops for 8 days and imaged. Two independent spheroids of L15 cells (0.5 x $10^6$) cultured in hanging drops for 7-(D,E), 14-(D',E'), or 21- (D",E") days and imaged. Scale bar: 100 μm.

**Supplementary Figure S6. L15-derived spheroids are mitotically active.** 7-day hanging drop culture derived L15 spheroids were fixed, immunostained and imaged. Immunostainings were performed for phosphoHistone3 (PH3, B) and DAPI (A). Merged image of PH3 and DAPI (C) is also shown. All images represent confocal projections of imaged Z series. Scale bar = 50 μm

**Supplementary Figure S7: Organization of cell junction markers in L15 spheroids.** 7-day hanging drop culture derived L15 spheroids were fixed, immunostained and imaged. Co-immunostainings were performed for Talin (A), DCAD (A', B'), and Discs large (Dlg, B). The surface exposed sections are represented in bluer tones and deeper sections in yellow tones. Each image is a projection of confocal sections This representation demonstrates that Talin is localized deeper than DCAD. However, similar analysis for Dlg and DCAD reveals that both are co-localized to sections at comparable depths. Scale bar = 50 μm

**Supplementary Figure S8. Expression of intestinal cell markers in L15 spheroids.** 7-day hanging drop culture derived L15 spheroids were fixed, immunostained and imaged. Immunostainings were performed for Horseradish peroxidase (HRP, C), Prospero (Pros, G), Nubbin (Nub, K) or Delta (Dl, O). The spheroids were co-stained with DAPI (A, E, I, M) and for GFP (B, F, J, N). Merged images of GFP and either HRP (D), Pros (H), Nub (L) or Dl (P) are also shown. Scale bar = 100 µm

**Supplementary Figure S9. Prospero is unevenly localized in L15 spheroids.** 7-day hanging drop culture derived L15 spheroids were fixed, immunostained and imaged. Two separate individual sections spaced five sections apart (A-D, surface exposed section and A’-D’, deeper section) are shown. Immunostainings were performed for Prospero (Pros, C, C’). The spheroids were also imaged for staining with DAPI (A, A’). Merged images of Pros and GFP (D, D’) are also shown. Scale bar = 100 µm.

### Supplementary Fig. S1

Stage *CG10116-GAL4; 20XUAS-6XGFP*

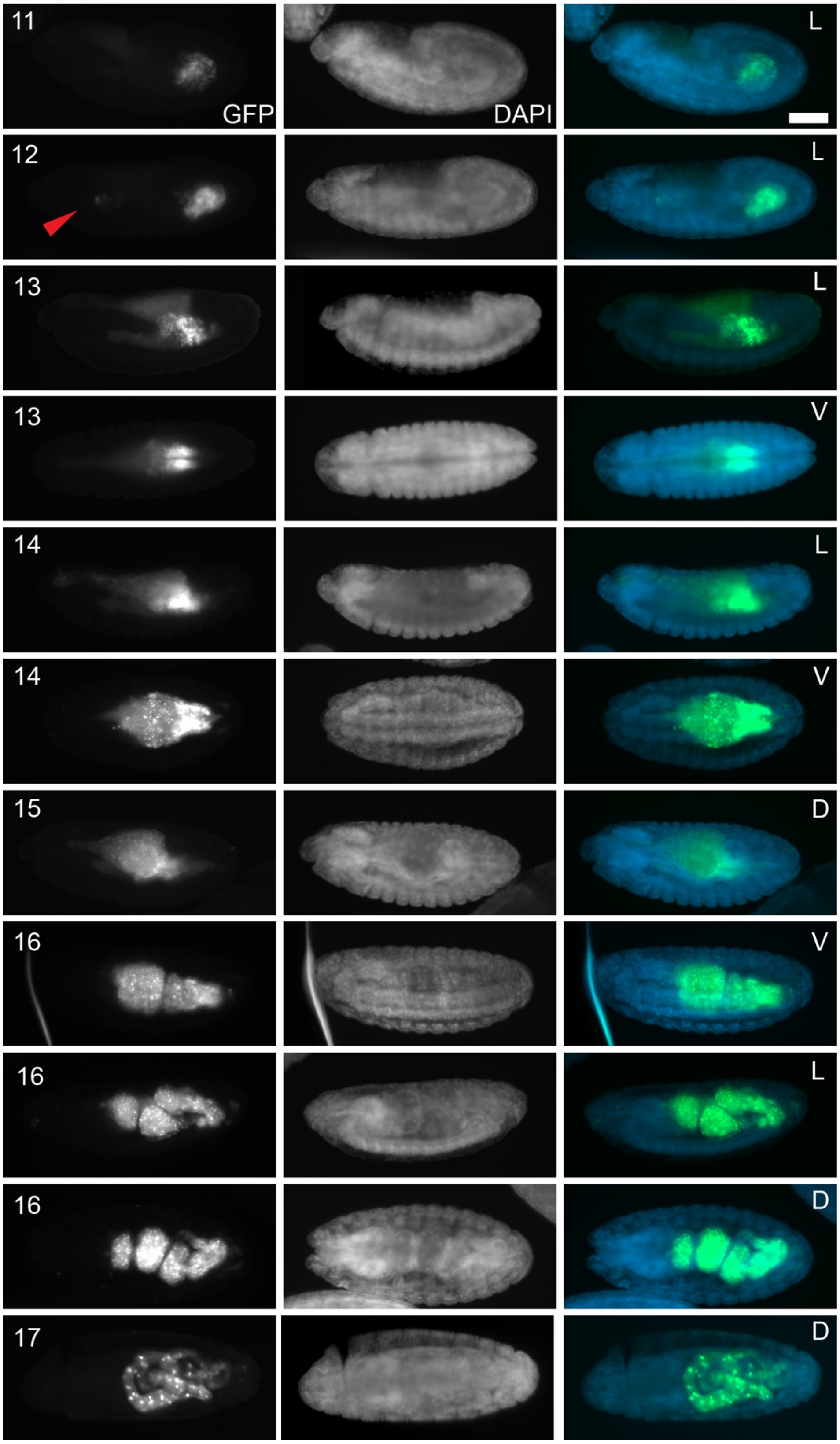

### Supplementary Fig. S2

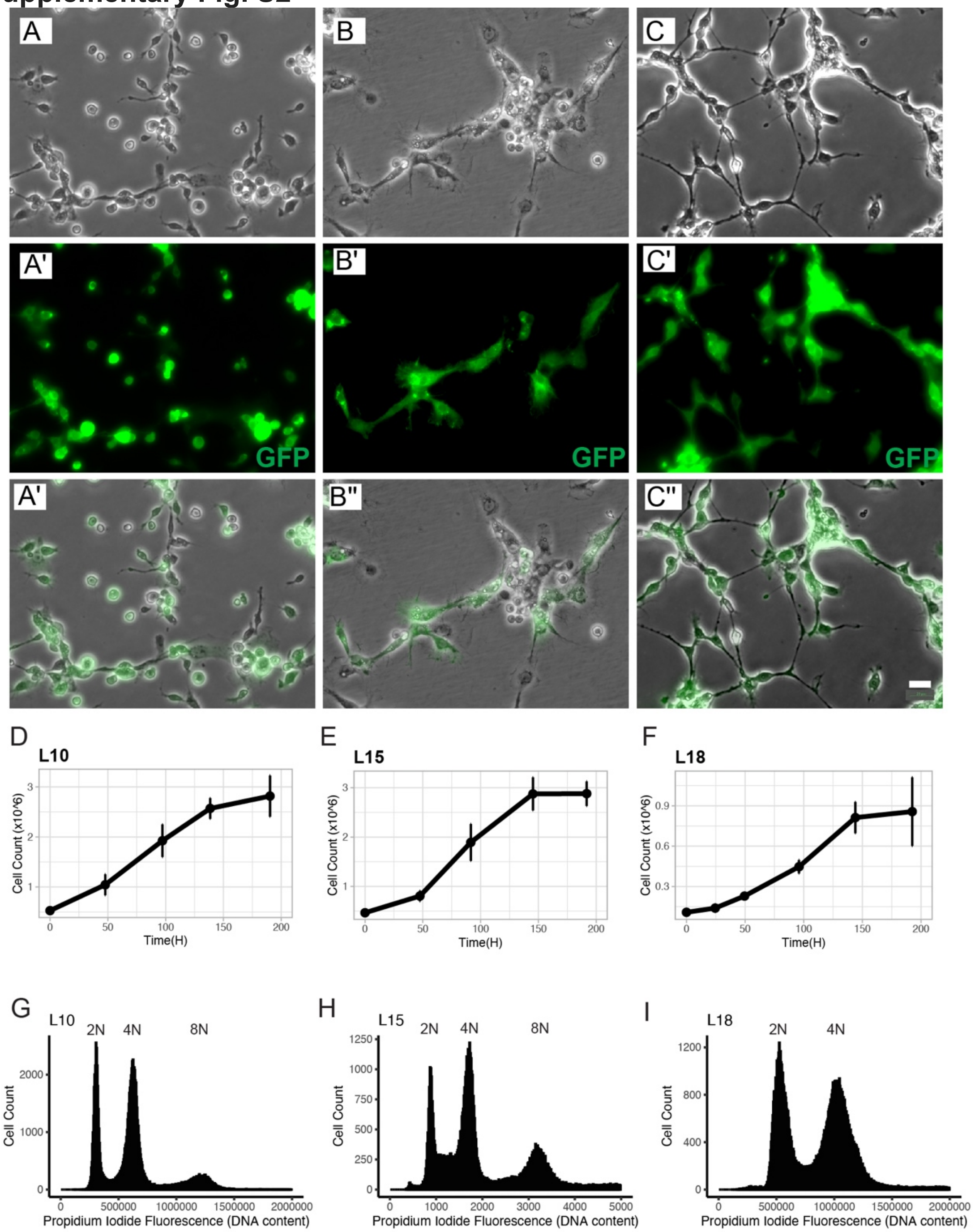


J

| Cell Line | Doubling Time (Hours) |
|---|---|
| **L10** | 69.81 |
| **L15** | 51.77 |
| **L18** | 53.25 |

**Supplementary Fig. S3**

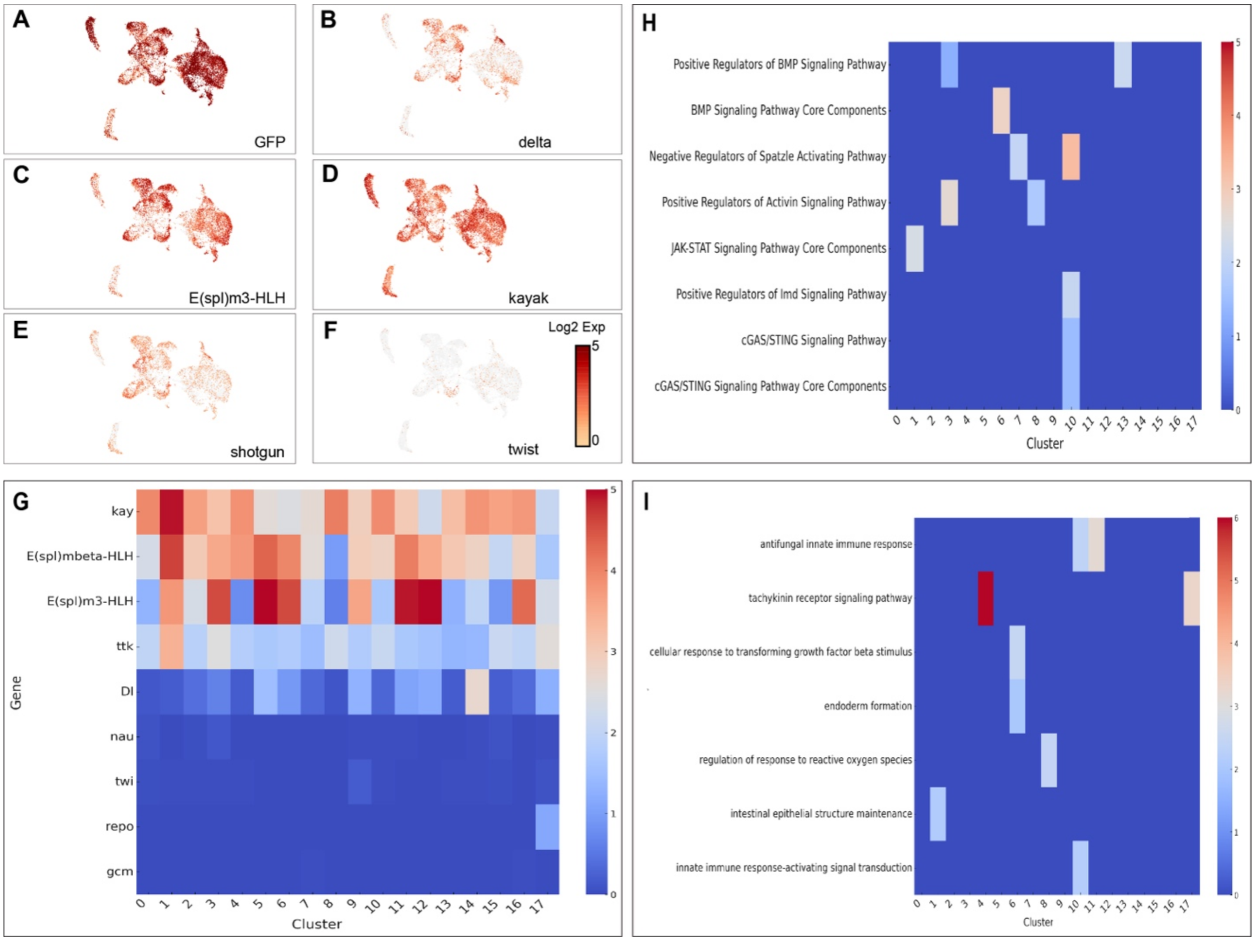

### Supplementary Fig. S4

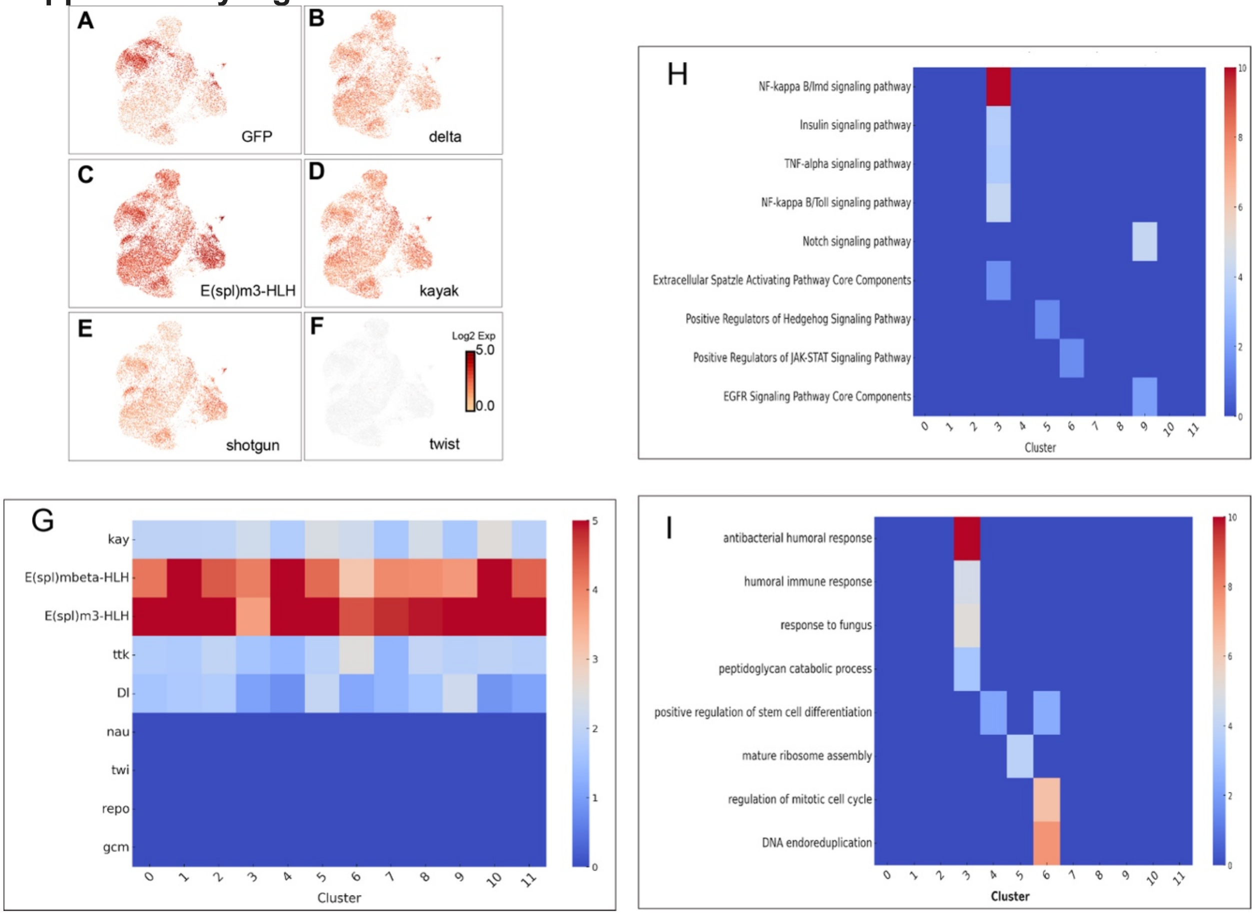

**Supplementary Fig. S5**

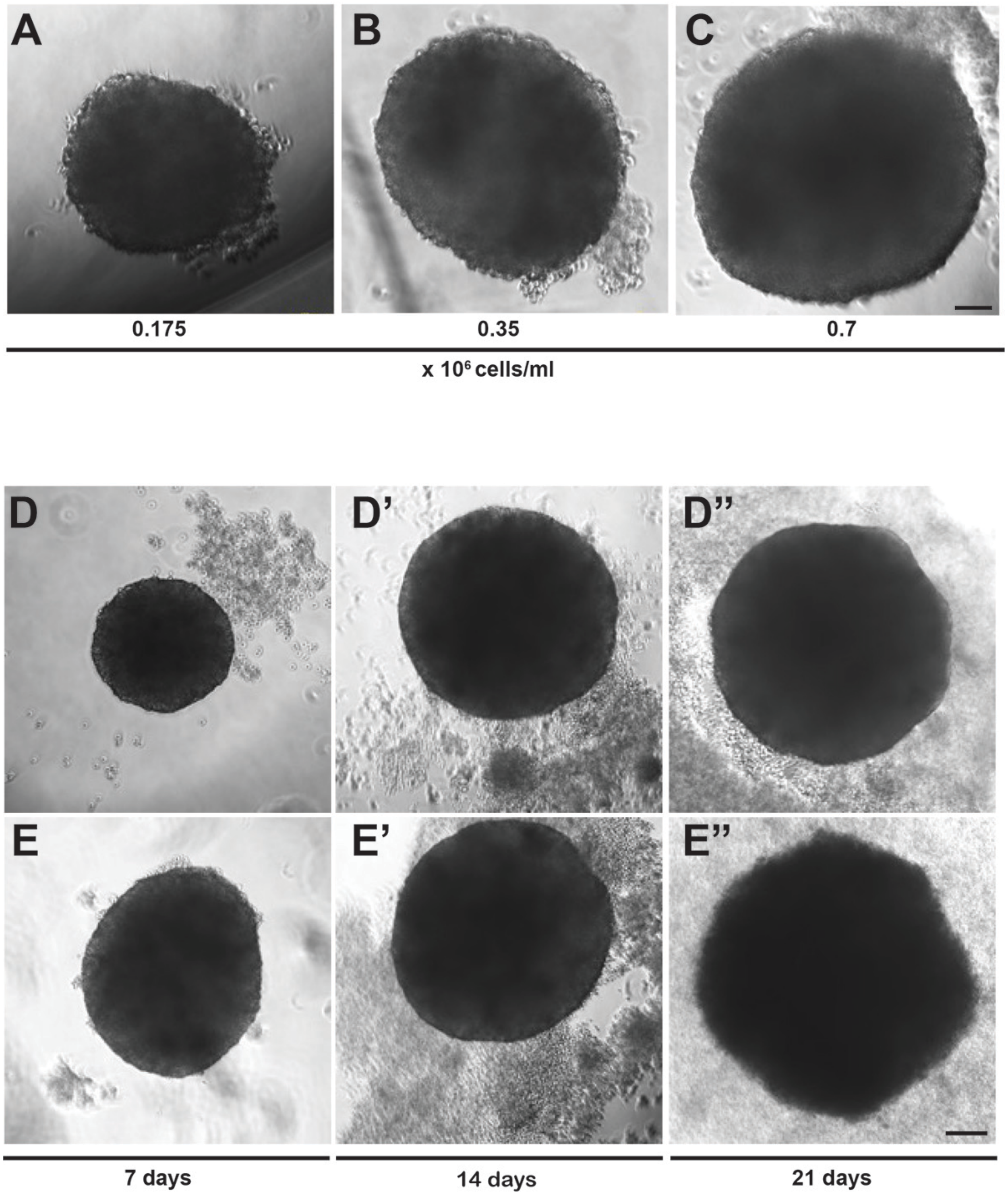

**Supplementary Fig. S6**

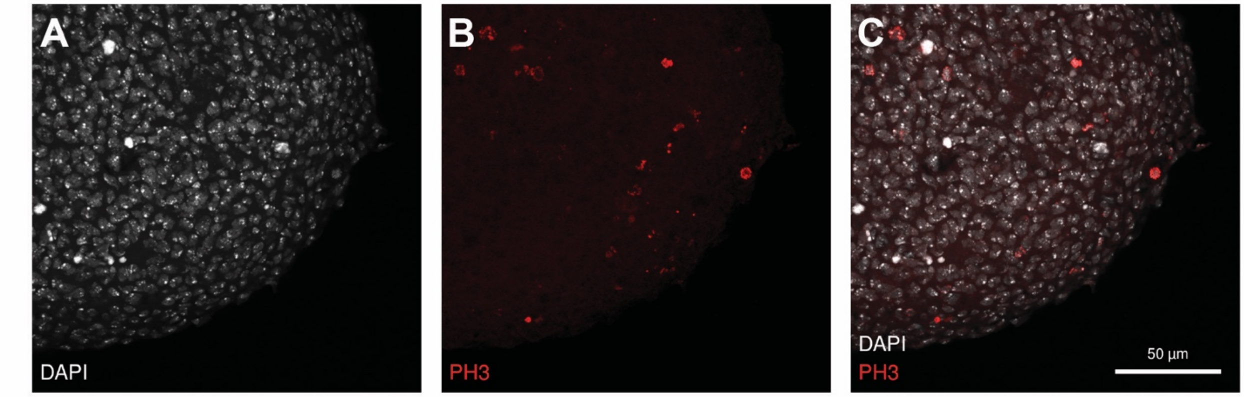

**Supplementary Fig. S7**

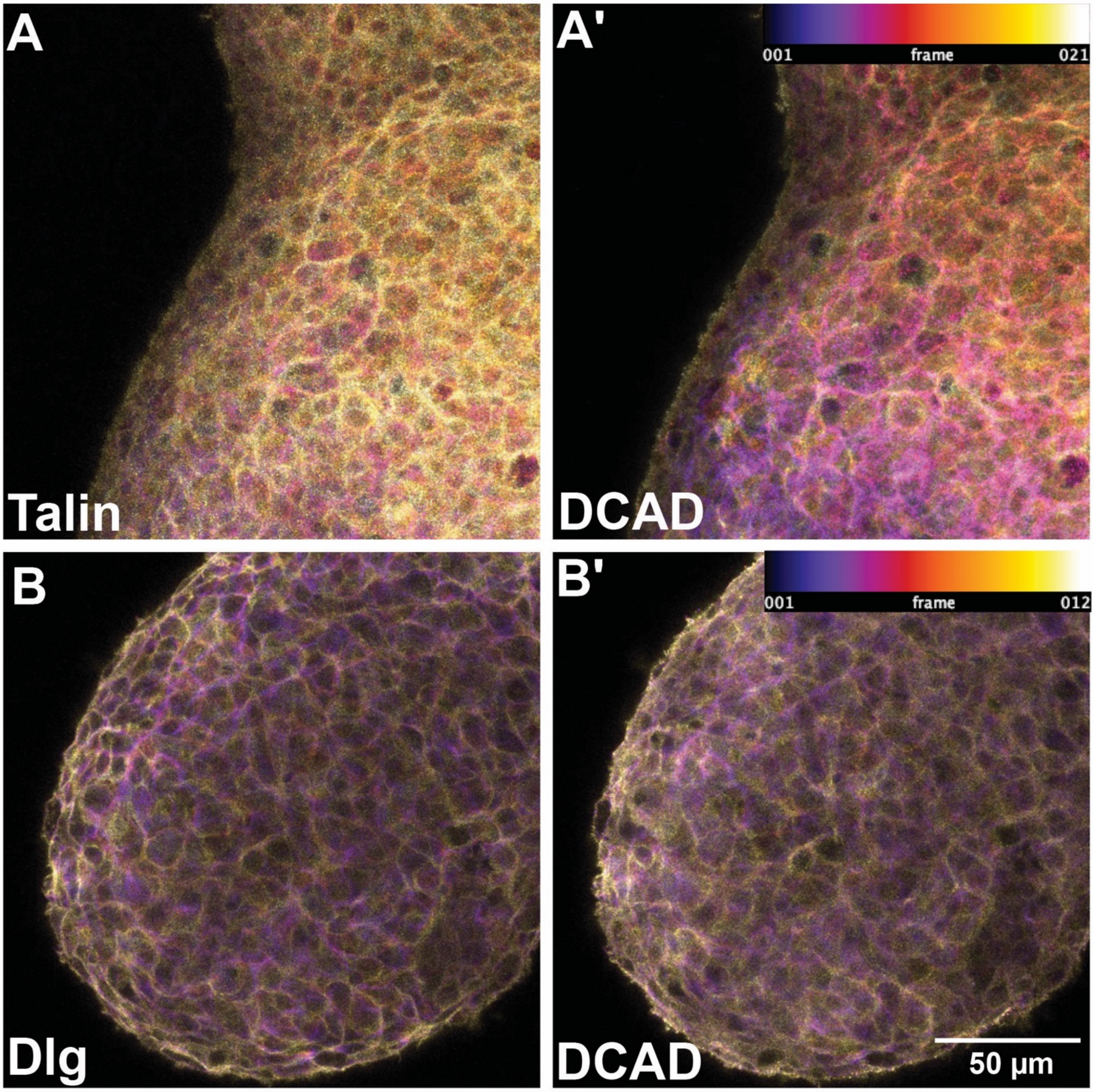


**Supplementary Fig. S8**

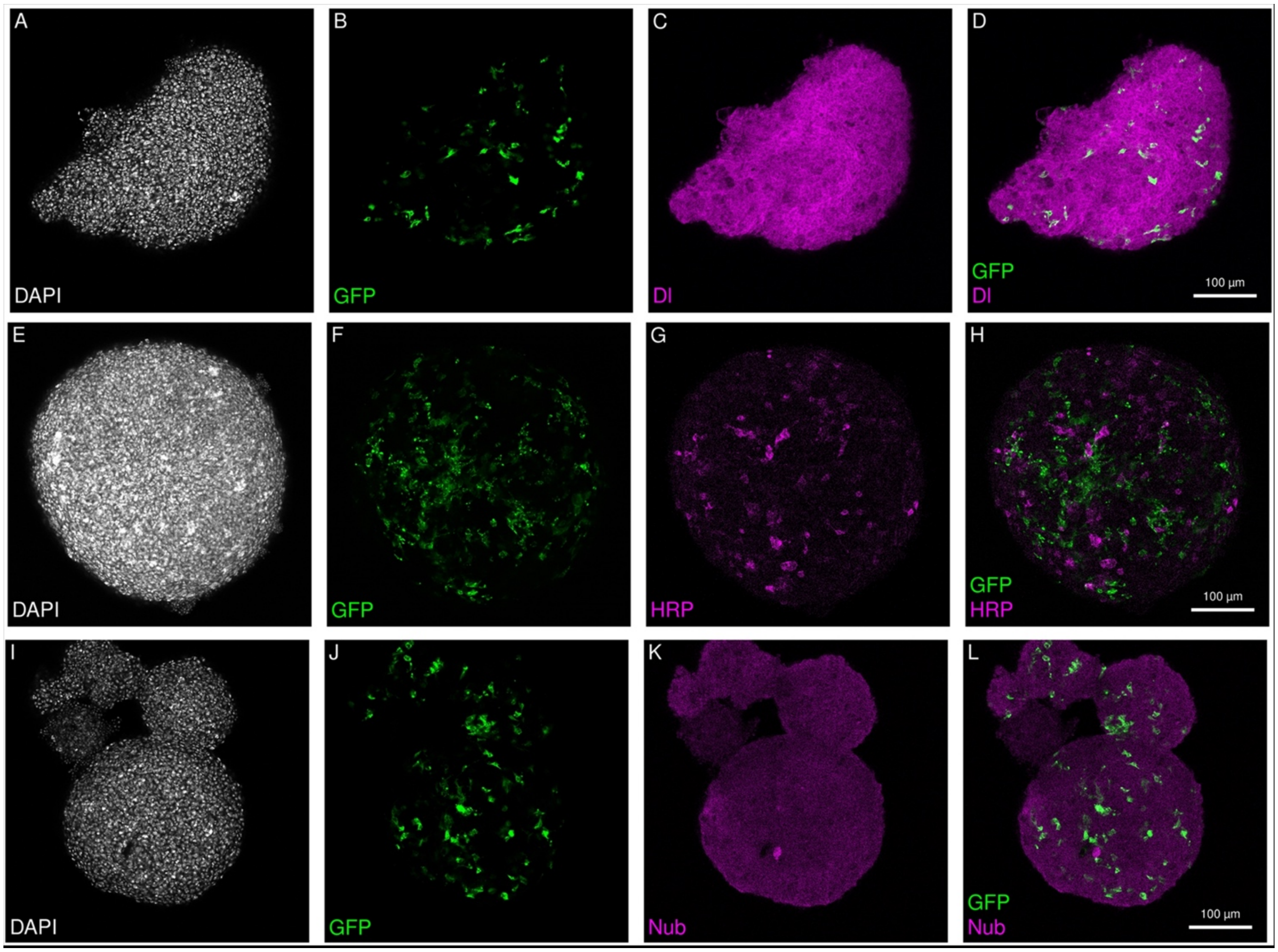
A
DAPI
B
GFP
C
Dl
D
GFP
Dl
100 µm
E
DAPI
F
GFP
G
HRP
H
GFP
HRP
100 µm
I
DAPI
J
GFP
K
Nub
L
GFP
Nub
100 µm
